# The Alzheimer risk factor CD2AP causes dysfunction of the brain vascular network

**DOI:** 10.1101/2020.12.10.419598

**Authors:** Milène Vandal, Adam Institoris, Ben Korin, Colin Gunn, Suzie Lee, Jiyeon Lee, Philippe Bourassa, Ramesh C. Mishra, Govind Peringod, Yulan Jiang, Sotaro Hirai, Camille Belzil, Louise Reveret, Cyntia Tremblay, Mada Hashem, Esteban Elias, Bill Meilandt, Oded Foreman, Meron Rouse-Girma, Daniel Muruve, Wilten Nicola, Jakob Körbelin, Jeff F. Dunn, Andrew P. Braun, David A. Bennett, Grant R.J. Gordon, Frédéric Calon, Andrey S. Shaw, Minh Dang Nguyen

## Abstract

Genetic variations in CD2-associated protein (CD2AP) predispose to Alzheimer’s disease (AD) but the underlying mechanisms remain unknown. Here, we show that a cerebrovascular loss of CD2AP is associated with cognitive decline in AD and that genetic downregulation of CD2AP in brain endothelial cells impairs memory function in two distinct mouse models. Mice with reduced CD2AP in brain microvessels display decreased resting cerebral blood flow, impaired functional hyperemia and vasomotion. In brain endothelial cells, CD2AP regulates the levels and signaling of ApoE receptor 2 elicited by Reelin glycoprotein. Activation of the CD2AP-ApoER2 pathway with Reelin mitigates the toxic effects of Aβ on resting blood flow and vasomotion of brain vessels depleted of CD2AP. Thus, we demonstrate that deregulation of CD2AP perturbs specific functions and segments of the cerebral microvasculature and propose that targeting CD2AP molecular partners may offer refined therapeutic strategies for the treatment of AD.

## Introduction

The brain vasculature is composed of a complex network of vessels with specialized segments presenting distinct cellular and molecular signatures (Schaeffer and Iadecola, 2021). Each type of vessels along the vascular tree plays unique and critical roles in regulating cerebral blood flow (CBF) that is essential to supply oxygen and nutrients to the brain. Cerebrovascular dysfunction that involves brain endothelial cells (BECs) and other cell types forming the neurovascular unit, is amongst the earliest defects observed in AD (Korte et al., 2020). Recent transcriptomic mapping studies of BECs reveal a remarkable heterogeneity across vascular segments (Schaeffer and Iadecola, 2021) that are differentially affected in AD patients (Yang et al., 2022). As such, the response to toxic Aβ species varies between pial vessels, arterioles and capillaries (Niwa et al., 2000; Nortley et al., 2019). Deciphering the molecular pathways underlying the dysfunction of specific brain vessel types could provide important clues for the treatment of AD.

Polymorphisms in the *CD2AP* gene represent one of the top 10 genetic predisposition factors for AD (alzgene.org) (Chen et al., 2015; Hollingworth et al., 2011). CD2AP (CD2-associated protein) encodes a scaffolding protein that participates to remodeling of the actin cytoskeleton and membrane trafficking during receptor endocytosis and cell-cell interactions (Kobayashi et al., 2004; Lynch et al., 2003). A few studies have linked CD2AP to amyloid precursor protein processing and tau-induced neurodegeneration (Ubelmann et al., 2017; Furusawa et al., 2019) but there is little (no) functional evidence relating AD pathophysiology and cognitive impairment. Interestingly, immunostaining data show that the protein is expressed in the brain vasculature (Li et al., 2000) and single-cell RNAseq reveal that CD2AP is enriched in BECs in both humans and mice (Lehtonen et al., 2008; Li et al., 2000; Vanlandewijck et al., 2018; Yang et al., 2022). Yet, how brain endothelial CD2AP contributes to AD remains a mystery.

To address this knowledge gap, we measured CD2AP protein levels in brain microvessels from AD volunteers and based on our findings showing reduction of CD2AP in the brain vascular fraction, we generated two mouse models with CD2AP genetically depleted in BECs. Using these models, we performed a stratified characterization of the roles of CD2AP in brain capillaries, penetrating arterioles and pial arteries to assess neural activity-dependent changes in red blood cell movements, microvessel diameter, and resting measurements of cerebral blood perfusion. We then identified a novel Reelin-ApoER2-CD2AP axis in BECs and discovered that endothelial CD2AP modulates the response to Reelin and Aβ in specific brain vessel types in vivo.

## Results

### Low levels of CD2AP in brain vessels are associated with cognitive dysfunction in AD volunteers

We first showed the enrichment of CD2AP protein in human brain vessel preparations by Western blot (Figure 1A) and immunostaining (Figure 1B), in line with previous mRNA findings in murine and human cerebral tissues (Lehtonen et al., 2008; Li et al., 2000; Vanlandewijck et al., 2018; Yang et al., 2022). To gain insights into the vascular role of CD2AP in AD pathogenesis, we compared vascular and total levels of CD2AP in parietal cortices of persons with no cognitive impairment (NCI) mild cognitive impairment (MCI) or a clinical diagnosis of Alzheimer’s disease (AD) from the Religious Order Study, a longitudinal clinical-pathologic study of aging and dementia (Bennett et al., 2006; Bourassa et al., 2019; Tremblay et al., 2017) (Table S1). Cerebrovascular concentrations of CD2AP were lower in AD compared to controls (Figure 1C) whereas levels in whole cortex homogenates remained similar between the groups (Figure 1D). No difference was seen in the levels of CD31 endothelial cell marker amongst the groups, suggesting that the lower levels of CD2AP in the AD group were not caused by a significant loss of BECs (Figure S1 and (Bourassa et al., 2020)). Cortical levels of CD2AP were inversely correlated with the amount of soluble Aβ42, Aβ40 and insoluble tau (Table S2), consistent with previous observations made in cell cultures (Ubelmann et al., 2017), *Drosophila melanogaster* (Shulman et al., 2014) and human CSF (Xue et al., 2021). Importantly, AD individuals with the worst cognitive function displayed the lowest levels of CD2AP in their post-mortem brain vascular fraction (Figure 1E and Table S2); this correlation that was not found for CD2AP levels in total cortex samples (Figure 1F). Overall, our data indicate that a reduction of CD2AP in brain vessels is associated with more severe cognitive decline in AD subjects.

**Figure 1:**
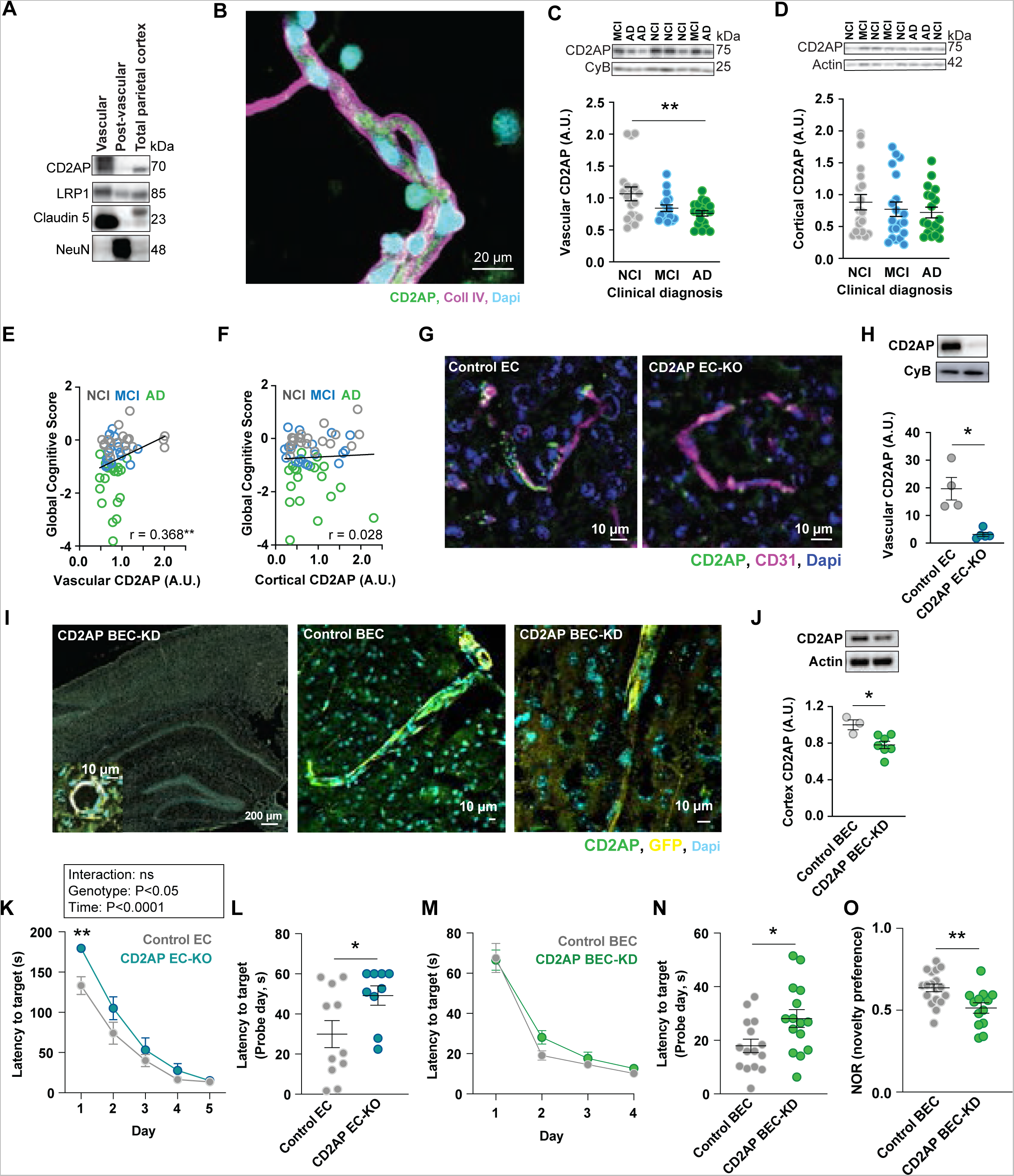
Lower levels of brain endothelial CD2AP are associated with cognitive dysfunction in AD. A) Enrichment of CD2AP in the cortex vascular fraction of human volunteers. Vascular = brain vessels enriched fraction, post-vascular = brain vessels depleted fraction, Total parietal cortex = unfractionated sample. B) Immunofluorescent image of a typical brain vessel isolated from a subject of the Religious Order Study cohort stained for CD2AP (white), Collagen IV (Coll IV, magenta) and Dapi (cyan). C) Vascular levels CD2AP in volunteers classified according to clinical diagnosis: no cognitive impairment (NCI, n = 19), mild cognitive impairments (MCI, n = 18) and Alzheimer’s disease (AD, n = 19). *Welch ANOVA followed by Dunnett’s T3 Post Hoc analysis adjusted for age and sex. Values are normalized on CyB levels. D) Cortical CD2AP levels in volunteers classified according to clinical diagnosis: no cognitive impairment (NCI, n = 19), mild cognitive impairments (MCI, n = 18) and Alzheimer’s disease (AD, n = 19). *One-way ANOVA followed by Tukey Post Hoc analysis. Values are normalized on actin levels. E) Correlation between the levels of vascular CD2AP and the global cognitive score. *Pearson r. adjusted for age and sex. F) Correlation between the levels of cortical CD2AP and the global cognitive score. *Pearson r. G) Immunofluorescent image of brain slice from Control EC and CD2AP EC-KO mice stained for CD2AP (green), CD31 (magenta) and Dapi (blue). H) Levels of vascular CD2AP in the brain of Control EC (n = 4 mice) and CD2AP EC-KO (n = 5 mice) mice quantified by Western blot (log from normalized value on CyB). *Unpaired Student t-test. I) Immunofluorescent image of brain slice from CD2AP BEC-KD and control mice stained for CD2AP (magenta), GFP (white) and Dapi (cyan). J) Levels of cortical CD2AP in the brain of Control BEC and CD2AP BEC-KD mice quantified by Western blot. (Control BEC: n = 3 mice, CD2AP BEC-KD: n = 7 mice). *Unpaired Student t-test. K) Learning curves for the time to reach the target during the Barnes Maze in Control EC and CD2AP EC-KO mice. *Repeated Measures ANOVA. L) Time to reach the target during the probe trial of the Barnes Maze in Control EC (n = 11 mice) and CD2AP EC-KO (n = 9 mice) mice. *Unpaired Student t-test. M) Learning curves for the time to reach the target during the Barnes Maze in Control BEC and CD2AP BEC-KD mice. *Repeated Measures ANOVA. N) Time to reach the target during the probe trial of the Barnes Maze in Control BEC (n = 15 mice) and CD2AP BEC-KD (n = 15 mice) mice. *Unpaired Student t-test. O) Novelty preference during the novel object recognition task in Control BEC (n = 18 mice) and CD2AP BEC-KO (n = 13 mice) mice. *Unpaired Student t-test. Data are presented as mean ± SEM. *P<0.05, **P<0.01.

### Loss of CD2AP in endothelial cells of mice impairs memory function

We next assessed whether loss of CD2AP in the brain endothelium has consequences on memory function in mice. Since CD2AP null mice die around 6 weeks of age from renal failure (Dustin et al., 1998; Shih et al., 1999), we generated two new mouse models. First, CD2AP endothelial cell-knockout (EC-KO) mice were obtained by crossing a *CD2AP^flox^*mouse with an endothelial-specific *Tie2-Cre* line (Kisanuki et al., 2001). Loss of CD2AP in brain vessels was confirmed by immunostaining (Figure 1G) and Western blot on vessels isolated from mouse brains (Figure 1H). Second, CD2AP brain endothelial cell-knocked down (BEC-KD) mice were generated by single retro-orbital injection of adeno-associated viruses (AAV) targeting specifically BECs (Körbelin et al., 2016; Nikolakopoulou et al., 2021; Yousef et al., 2019). Viral vectors contained short hairpin RNAs (shRNAs) directed against CD2AP mRNA or a random non-specific sequence (Control). CD2AP siRNAs caused a 70-80% selective decrease of CD2AP in primary mouse BECs and in a human BEC line, confirming their specificity (Figure S2). In mice, the AAVs infected BECs lining blood vessels, visualized with immunostaining (Figure 1I and Figure S2). After three weeks, a timeline known for the viral vector to be expressed (Körbelin et al., 2016), CD2AP BEC-KD mice displayed a significant knockdown of CD2AP protein levels in the cortex when compared to mice injected with the control viral vector (control mice), as demonstrated by Western blots (Figure 1J). No infection of mural cells, astrocytes, and limited infection of neuronal cells was observed (Figure S2).

CD2AP EC-KO mice showed no overt phenotype when compared with control animals, moving, and feeding normally up to the age of 15-month-old, before sacrifice for experimental procedures. Since CD2AP is important for kidney function (Dustin et al., 1998; Shih et al., 1999), and the protein is expressed in the renal vasculature (Jourde-Chiche et al., 2019), we evaluated kidney morphology and immune cell populations in CD2AP EC-KO mice. Compared to controls, CD2AP EC-KO mice did not display any gross renal pathologic changes or differences in resident immune cell composition (Figure S3). Next, we evaluated CD2AP EC-KO and CD2AP BEC-KD mice for spatial learning and memory with the Morris Water Maze and the Barnes Maze tasks, working memory with the novel object recognition tests, locomotion and anxiety-like behaviors using the open field test. None of the mouse models displayed anxiety-like behaviors and locomotion deficiency as indicated by the distance travelled and the center/periphery ratio in the open field test (Figure S4). In the Morris Water Maze task, 6-month-old CD2AP EC-KO mice took more time to reach the visible platform on the 3^rd^ day when compared to control mice (Figure S4). At 12 months of age, the mutant mice showed slower learning (Figure 1K) and worse memory (Figure 1L) than control mice during the Barnes Maze. In the novel object recognition, CD2AP EC-KO mice showed the same level of interest toward the novel object than control mice (Figure S4). During the Barnes Maze, CD2AP BEC-KD mice had similar learning than control mice (Figure 1M) and took more time to find the target during the probe trial (Figure 1N). However, they displayed lower recognition index in the novel object recognition test (Figure 1O), indicative of impaired working memory function. Thus, the two mouse models are distinct in targeting endothelial CD2AP (acute depletion in adult brain endothelial cells vs depletion at birth in the whole body-endothelium) but they highlight the common finding that genetic downregulation of CD2AP the protein in endothelial cells alters memory function.

### Brain endothelial CD2AP regulates resting cerebral blood flow and neurovascular coupling throughout the vascular network

Memory loss can be associated with brain hypoperfusion and defects in processes that control cerebrovascular function such as neurovascular coupling (Millar et al., 2020; Grady and Garrett, 2014). As both CD2AP mutant mouse models displayed cognitive deficits and CD2AP is a candidate for the lower resting CBF observed in AD patients (Yao et al., 2019), we evaluated resting CBF and neurovascular coupling in our mice. Using arterial spin labelling magnetic resonance imaging (MRI), we found that both CD2AP EC-KO and CD2AP BEC-KD mice exhibited a modest reduction in resting CBF over time (Figure 2A to D). Overall, in both mouse lines, genotype had no impact on resting CBF (Figure 2A and B). However, when looking separately in Controls and CD2AP EC-KD mice, resting CBF decreased in the hippocampus of CD2AP EC-KO mice as the mice aged from 6 to 14 months (Figure 2A, D and S5). CD2AP BEC-KD also displayed reduced resting CBF in the cortex between the second and the third week following AAV injection (Figure 2B and D). Such progressive reduction was not observed in animals injected with the control virus (Figure 2A, B and D). Furthermore, we found a positive correlation between cortical CBF and CD2AP levels in CD2AP BEC-KD animals: lower levels of CD2AP were associated with lower resting CBF (Figure 2C). Thus, endothelial CD2A*P has a modest yet significant impact on resting CBF in an age-dependent manner. Since isoflurane is a potent vasodilator (Lyons et al., 2016; Tran and Gordon, 2015), the anesthetic might interfere with CBF measurement during MRI. We therefore performed awake mouse two-photon fluorescence microscopy to investigate vessel function (Figure 2E). We confirmed that the AAVs infected the brain vasculature (Figure 2F). The fluorescent signal was detected in pial arteries, penetrating arterioles and capillaries and no obvious morphological differences were observed in CD2AP BEC-KD compared to control mice. We first measured capillary red blood cells flux, a proxy to resting CBF (Kleinfeld et al., 1998)(Figure 2G) and found that CD2AP BEC-KD mice had a lower resting capillary flux compared to controls (Figure 2H). We next assessed neurovascular coupling by measuring functional hyperemia in the barrel cortex of awake CD2AP BEC-KD and control mice following stimulation of facial whiskers (Tran et al., 2018). The propagation and intensity of this sensory-induced blood flow in response to changes in regional neuronal activity is controlled in part by the vascular endothelium at the levels of capillaries (Longden et al., 2021), penetrating arterioles (Chow et al., 2020), and pial arteries (Chen et al., 2014). In CD2AP BEC-KD, the maximum speed of brain capillary red blood cells was ∼40% lower (Figure 2I to L) when compared to that of control mice during functional hyperemia. The time to reach maximum red blood cell speed in response to whisker stimulation in brain capillaries was also delayed in the mutant mice (Figure 2K). Similarly, the maximal vessel dilation in response to whisker stimulation was reduced by ∼40% in penetrating arterioles (Figure 2M and N) and by ∼30% in pial arteries (Figure 2O and P) of CD2AP BEC-KD mice. Together these results show that CD2AP reduction impairs blood flow regulation in different segments of the vascular network at resting state and during enhanced neuronal activity.

**Figure 2:**
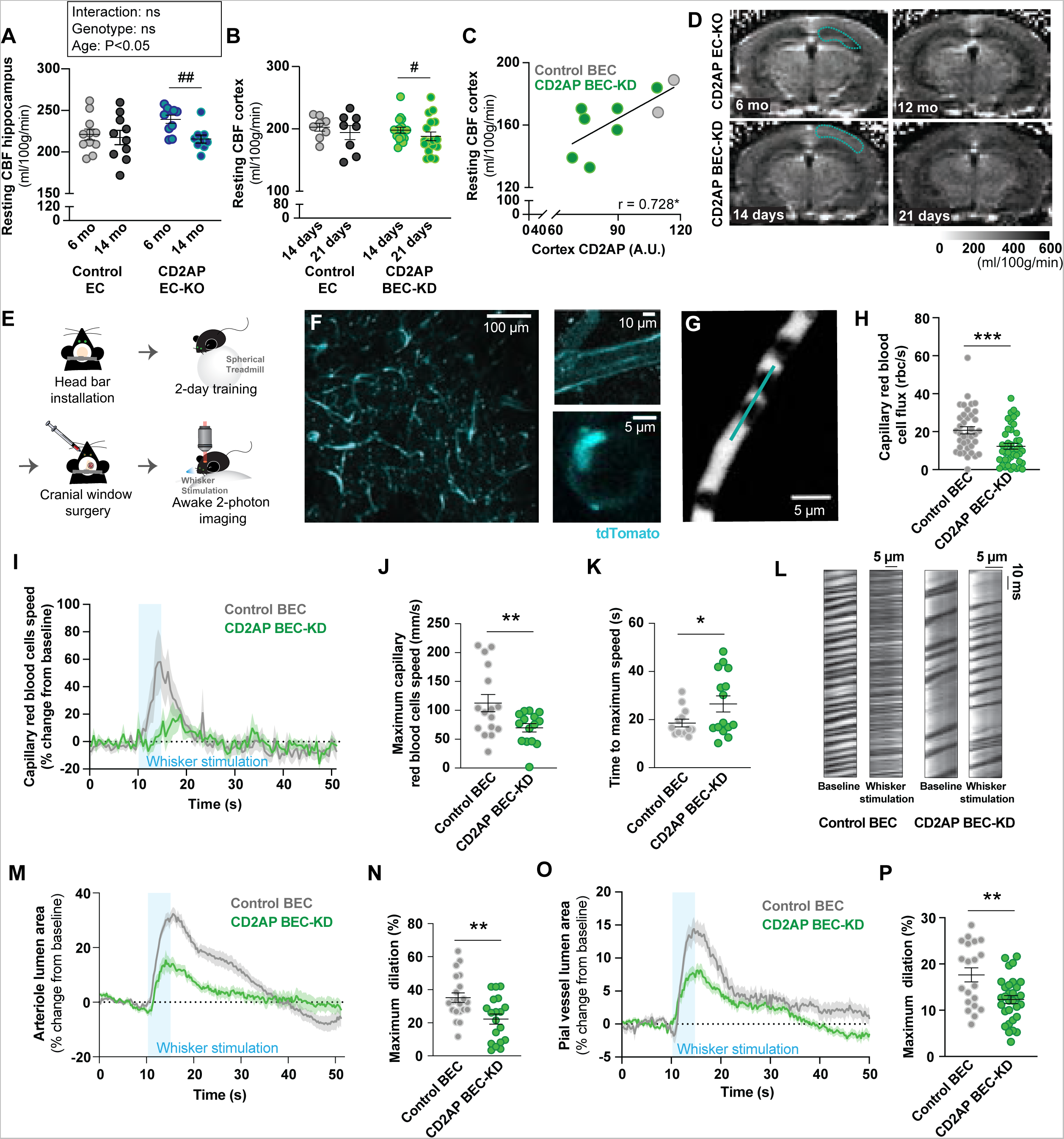
Endothelial CD2AP regulates resting CBF and neurovascular coupling. A) Resting hippocampus CBF in Control EC and CD2AP EC-KO mice at 6 months (Control EC: n = 11 mice, CD2AP EC-KO: n = 10 mice) and 14 months (Control EC: n = 10 mice, CD2AP EC-KO n = 9 mice) of age. *Two-way ANOVA, ^#^Unpaired Student t-test. B) Resting cortex CBF in Control BEC (n = 9 mice) and CD2AP BEC-KD (n = 17 mice) mice 14 and 21 days after viral vector injection. *Repeated Measures ANOVA, ^#^Paired Student t-test. C) Correlation between resting cortex CBF and CD2AP levels, 21 days after AAV injections. *Pearson r. D) Representative images of brain perfusion in CD2AP EC-KO and CD2AP BEC-KD mice taken during MRI. The cyan line represents the zone analyzed for CBF. E) Illustration representing the awake two-photon imaging experimentation setup. F) Representative image of the brain vasculature of a mouse injected with the AAVBR1 virus on the awake two-photon microscope. Cyan = tdTomato. G) Example of a capillary selected for line scan imaging. H) Resting capillary red blood cells flux in Control BEC (14 mice, 38 recordings) and CD2AP BEC-KD (14 mice, 43 recordings) mice. Each dot represents one recording. *Unpaired Student t-test. rbc = red blood cell. I) Change in capillary red blood cells speed in response to whisker stimulation (5 s) in Control BEC (5 mice, 16 stimulations) and CD2AP BEC-KD (6 mice, 15 stimulations) mice. Each value represents the average of 4 time points. J) Maximal capillary red blood cells speed after whisker stimulation. Each dot represents one stimulation. *Unpaired Student t-test with Welch correction. K) Time to reach the maximum red blood cells speed after whisker stimulation. Each dot represents one stimulation. *Unpaired Student t-test with Welch correction. L) Representative example of line scan from capillary of Control BEC and CD2AP BEC-KD mice before and after whisker stimulation. M) Change in penetrating arterioles luminal area in response to whisker stimulation (5 s) in Control BEC (5 mice, 7 arterioles, 21 stimulations) and CD2AP BEC-KD (6 mice, 7 arterioles, 19 stimulations) mice. N) Penetrating arterioles maximal dilation after whisker stimulation. Each dot represents one stimulation. *Unpaired Student t-test. O) Change in pial vessels luminal area in response to whisker stimulation (5 s) in BEC-Control (9 mice, 9 vessels, 20 stimulations) and CD2AP BEC-KD (9 mice, 11 vessels, 31 stimulations) mice. P) Maximal pial vessels dilation after whisker stimulation. Each dot represents one stimulation. *Unpaired Student t-test. Data are presented as mean ± SEM. ^#^P<0.05, ^##^P<0.01, *P<0.05, **P<0.01.

### CD2AP controls ApoER2 levels and Reelin-mediated signaling in BECs

To gain insights into the mechanisms by which CD2AP regulates cerebrovascular function through the endothelium, we took a candidate approach to find functional partners for CD2AP. The trafficking of ApoER2, the highest expressed ApoE receptor in BECs (Vanlandewijck et al., 2018), is regulated by the CD2AP ortholog Cbl interacting protein 85 (Fuchigami et al., 2013). Therefore, we hypothesized that ApoER2 is a target for CD2AP in the brain endothelium. We first confirmed the enrichment of CD2AP in cultured mouse primary BECs using Western blot (Figure 3A) and immunostaining (Figure 3B). Next, using a human BEC line (hCMEC/D3) and mouse brain tissues, we tested the potential association between CD2AP and ApoER2. We found that ApoER2 co-immunoprecipitates with CD2AP and this association was detectable in lysis buffers containing a range of concentrations of ionic and non-ionic detergents (Figure 3C to E). Dynamin 2, a key protein for receptor endocytosis and membrane trafficking, was also detected in CD2AP immunoprecipitates (Figure 3E), supporting a role for CD2AP in ApoER2 trafficking and consistent with the known role for CD2AP in these cellular processes (Kobayashi et al., 2004). No interaction was found between CD2AP and PICALM (Phosphatidylinositol Binding Clathrin Assembly Protein, see Figure 3E), another endocytic molecule associated with increased risk for AD (Naj et al., 2014). In brain lysates from adult mice, CD2AP antibodies immunoprecipitated both CD2AP and ApoER2 (Figure 3D). The reverse co-immunoprecipitation using ApoER2 antibodies also pulled down CD2AP (Figure 3D).

**Figure 3:**
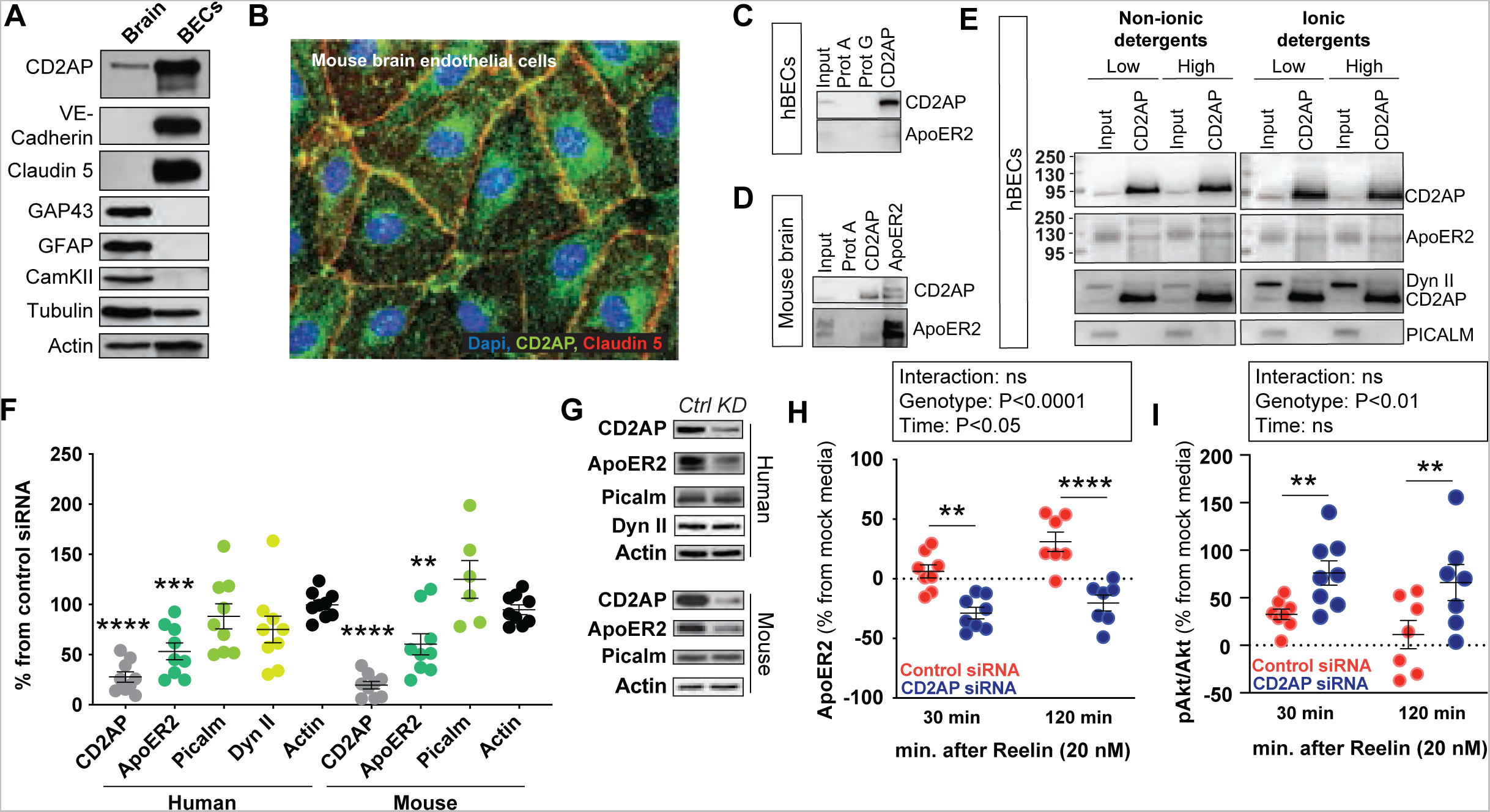
CD2AP regulates Reelin-mediated ApoER2 signaling and levels in brain endothelial cells. A) Western blot showing the enrichment of CD2AP in BECs compared to whole brain extracts. B) Immunohistochemistry for CD2AP in mouse primary BECs. Green: CD2AP, Red: Claudin 5, Blue: Dapi. C-E) Co-immunoprecipitation of CD2AP with ApoER2 in C) human brain endothelial cell line (hCMEC/D3, hBECs), D) adult mouse brain and E) hBECs under ionic and non-ionic detergent. F) Effect of CD2AP depletion on the levels of specific proteins in hBECs and mouse primary brain endothelial cells (mBECs). Each dot represents a different well. Data are presented as a % of the mean of cells transfected with control siRNA. *One sample t-test compared to 100. G) Representative Western blot images from the CD2AP and control siRNA transfection in hBECs and mBECs. H) ApoER2 and I) pAkt/Akt levels in hBECs in control or CD2AP KD cells treated with Reelin (20 nM) or control solution for 30 min and 120 min. Control and CD2AP KD 30 min (n= 8 wells), Control and CD2AP KD 120 min (n = 7 wells). Each dot represents a different well. *Two-way ANOVA followed by Sidak’s multiple comparisons test. **P<0.01, ***P<0.001, ****P<0.0001

To determine the role of CD2AP on ApoER2 function, we silenced the expression of *CD2AP* in mouse and human BECs using siRNA. Knockdown of CD2AP had no effect on the levels of PICALM or Dynamin II but caused a significant ∼40% decrease in levels of ApoER2 (Figure 3F and G). To investigate ApoER2 signaling in the absence of CD2AP in BECs, we treated CD2AP-depleted BECs with the glycoprotein Reelin, a well characterized ligand for ApoER2. Reelin mediates neuronal plasticity (Wasser and Herz, 2017) and confers protection against parenchymal Aβ toxicity in animal models of AD (Lane-Donovan et al., 2015; Pujadas et al., 2014) but its function in the brain vasculature remains largely undefined. The glycoprotein targets alpha-integrin, VLDLr and ApoER2 receptors that are expressed in the endothelium (Bock and May, 2016), ApoER2 being highly enriched in BECs by comparison to VLDLr (Vanlandewijck et al., 2018). In the canonical Reelin signaling pathway, the binding of Reelin to ApoER2/VLDLr triggers the phosphorylation of the adaptor protein disabled-1 (Dab1), and subsequently phosphorylation/activation of protein kinase B (Akt) at Ser473 (Beffert et al., 2002). Like untreated BECs lacking CD2AP (Figure 3F and G), CD2AP-depleted BECs treated with Reelin displayed lower levels of ApoER2 at 30 and 120 min post-treatment (Figure 3H). Akt activation (measured by p-Ser 473 Akt/total Akt ratio) was higher in CD2AP-depleted BECS at both time points when compared to control cells (Figure 3I), possibly to compensate for the decrease in ApoER2 levels. In brief, CD2AP functional interacts with ApoER2; the loss of CD2AP alters the levels of ApoER2 and deregulates Reelin-mediated signaling in BECs.

### Endothelial CD2AP controls Reelin-mediated signaling *in vivo*

Building on these findings, we used Reelin to investigate ApoER2 signaling *in vivo* in CD2AP BEC-KD and control mice. To determine specifically the effect of Reelin on brain vessels we freshly isolated single cerebral arteries from mouse brains using pressure myography (Figure 4A) and applied Reelin on them (passive diameter 146 ± 15 μm, n=12). The arteries were maintained at the pressure of 70 mmHg and modulated with vasoconstrictive (i.e. phenylephrine, 250 nM) or vasodilatory drugs (i.e. KCa channel activator SKA-31 (3 and 15 μM), bradykinin (0.5 μM), sodium nitroprusside (SNP 10 μM), Pinacidil (10 μM)) to confirm their overall health and functionality before and after the application of Reelin or control solution (Figure 4B and C). The preparation contains BECs and smooth muscle cells with no neuron or astrocyte. Bath application of Reelin (1 μM) consistently promoted vasodilation of small arteries whereas the control solution had no effect (Figure 4D). To isolate the effects of Reelin on BECs, we subjected endothelium-denuded arteries to the same drug treatments. In the absence of the endothelium, the vasodilatory effect of Reelin was reduced by ∼70% (Figure 4D). Taken together, our results indicate that Reelin can exert vasodilatory effect by targeting receptors expressed by the brain endothelium, and independently from receptors expressed on mural cells, astrocytes and neurons.

**Figure 4:**
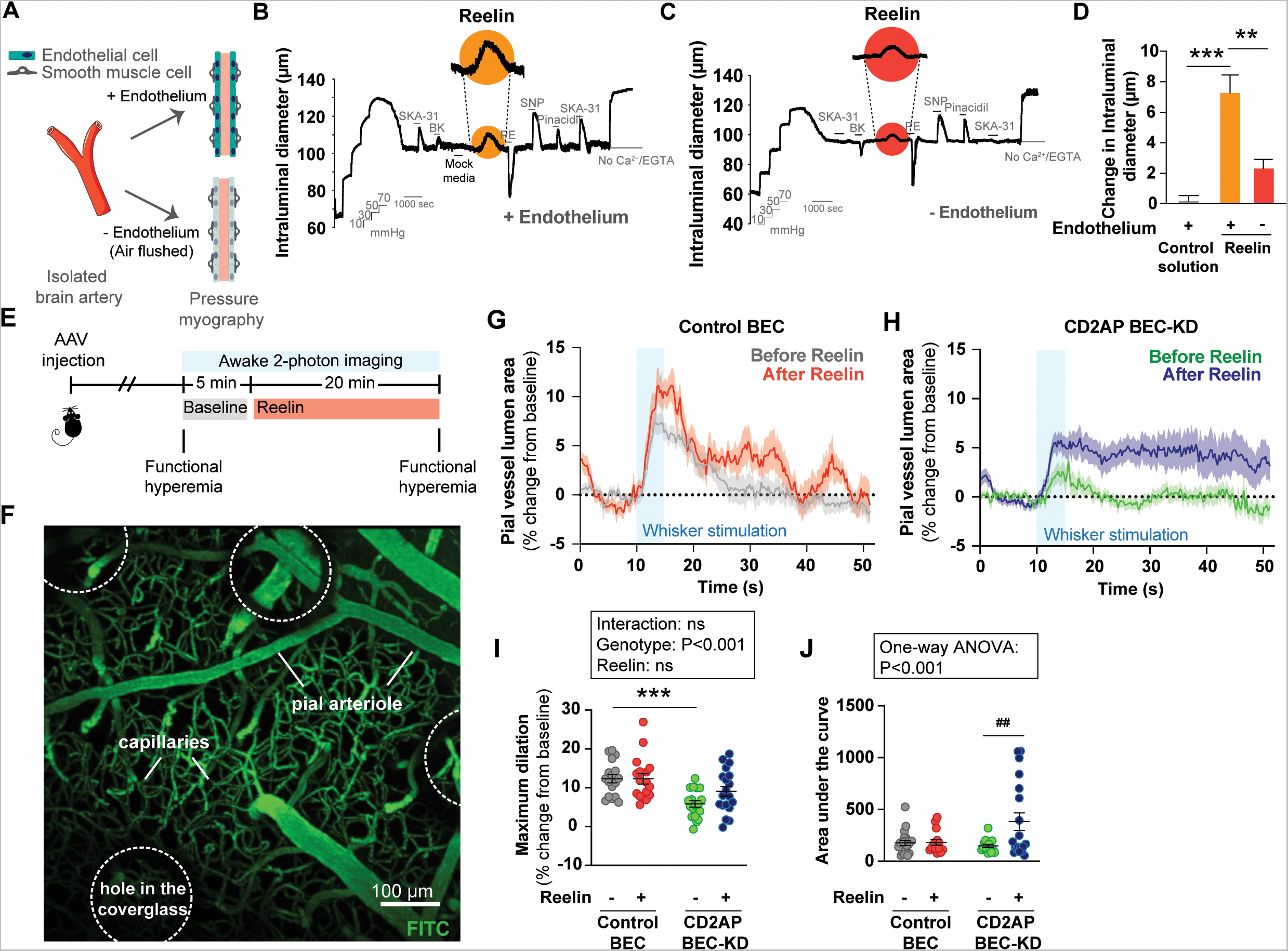
CD2AP regulates Reelin effect on brain vessels. A) Illustration representing the pressure myography experiment. Representative tracings displaying acute vasoactive responses to endothelium B) dependent and C) independent agents in single cannulated cerebral arteries from C57BL6 mice. SKA-31 (3 and 15 µM), bradykinin (BK, 0.5 µM), control solution (volume matched with Reelin), Reelin (1 µM), and the endothelium-independent agent phenylephrine (PE, 250 nM), sodium nitroprusside (SNP, 10 µM) and pinacidil (10 µM). Horizontal bars above the tracing indicate the treatment with each agent. D) Changes in intraluminal diameter of cerebral arteries triggered by the indicated vasoactive agents (n = 4 male mice). *One-way ANOVA followed by Tukey Post Hoc analysis. E) Timeline of the drug application during the experiment. F) Z-stack reconstructed image of the barrel cortex in a C57BL6 mouse mounted with a cover glass with holes and depicting FITC-dextran (green) vessels. Pial vessel transversal luminal area in response to whisker stimulation (5 s) in G) Control BEC (5 mice, 17 stimulations) and H) CD2AP BEC-KD (6 mice, 19 stimulations) infected mice before and during Reelin (1 μM) superfusion. I) Maximal dilation after whisker stimulation of pial vessels before and during Reelin. Each dot represents one stimulation. *Two-way ANOVA followed by Sidak’s multiple comparison test. J) Overall response to whisker stimulation. Each dot represents one stimulation. *One-way ANOVA. ^#^Unpaired t-test with Welch correction. Data are presented as mean ± SEM. **P<0.01, ***P<0.00, ^##^P<0.01

We next analyzed the effects of purified Reelin on the brain vessel tone *in vivo.* A solution of Reelin (1 μM) or a control solution was superfused onto the brain surface (Figure S6) (Tran and Gordon, 2015). Reelin consistently caused a robust dilation of penetrating arterioles (Figure S6), in line with our observation in *ex vivo* brain vessels preparation (Figure 4D). We next tested whether endothelial CD2AP modulates these functions of Reelin *in vivo*. We superfused Reelin for 20 min on the brain surface of awake CD2AP BEC-KD and control mice during two-photon imaging (Figure 4E and F). The effects of Reelin on pial vessel dilation in CD2AP BEC-KD and control mice was rather weak or subtle (see supplementary Figure S7). In the mutant mice, the vessels failed to return to baseline following the peak dilation in the presence of Reelin. As a result, the overall response to whisker stimulation was increased in the mutants after Reelin superfusion whereas Reelin was unable to enhance functional hyperemia in the controls (Figure 4J). Collectively these data suggest Reelin vasodilates in vivo but a positive influence on functional hyperemia is unmasked only when CD2AP in the brain endothelium is lowered.

### Loss of endothelial CD2AP increases A**β**40 accumulation and sensitizes specific vessels to A**β**42 oligomers

Cerebrovascular defects worsen Aβ accumulation while Aβ reduces CBF and impairs neurovascular coupling (Kisler et al., 2017). Since loss of CD2AP in the endothelium is linked to cerebrovascular dysfunction, we first investigated whether the loss of endothelial CD2AP impact Aβ accumulation. We crossed the PS2APP mouse model of brain β-amyloidosis (Ozmen et al., 2009) with CD2AP EC-KO mice (PS2APP CD2AP EC-KO) (Figure 5A). Overall, we observed a mild effect on Aβ accumulation in the PS2APP lacking endothelial CD2AP. No change was observed for soluble and insoluble Aβ42 accumulation in the cortex and hippocampus (Figure S8), in line with previous studies showing minimal effects of CD2AP on Aβ accumulation (Liao et al., 2015). However, loss of endothelial CD2AP increased insoluble Aβ40 in the hippocampus of PS2APP mice (Figure 5B to D). Aβ40 is the dominant species of Aβ that accumulates in cerebral amyloid angiopathy, commonly observed in AD (Bourassa et al., 2019), thereby suggesting that loss of endothelial CD2AP promotes vascular Aβ pathology.

**Figure 5:**
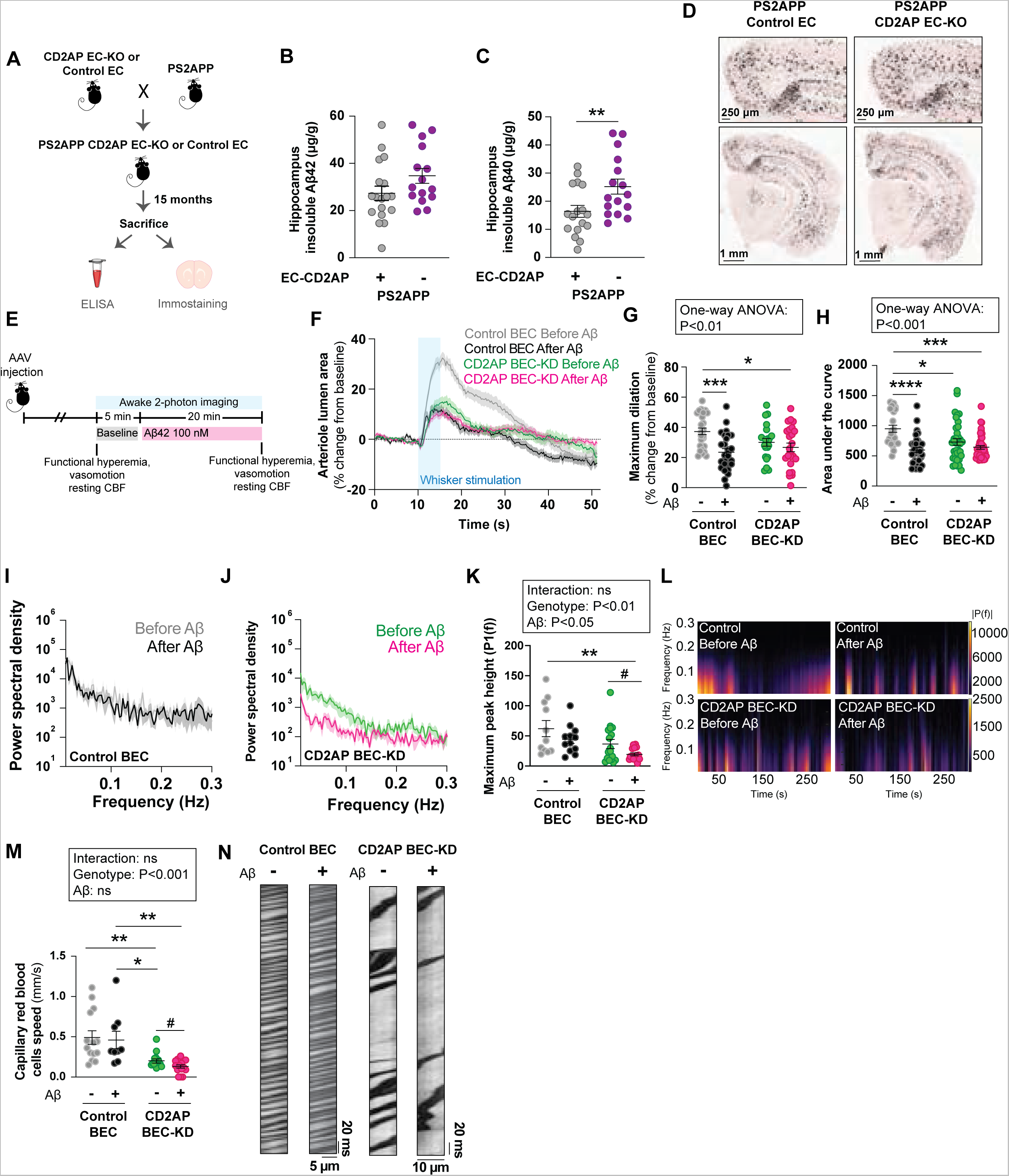
BECs CD2AP modulates capillary and arterioles response to A**β**. A) Representation of the PS2APP Control EC and CD2AP EC-KO mouse model. Levels of insoluble B) Aβ42 and C) Aβ40 in the hippocampus of PS2APP mice with (n = 17 mice) or without CD2AP (n = 16 mice) in the endothelium. Aβ were quantified by ELISA. *Unpaired Student t-test. D) Representative images of the Aβ40 staining in the PS2APP Control EC and CD2AP EC-KO. E) Timeline of the drug application during the experiments. F) Penetrating arterioles transversal luminal area in response to whisker stimulation (5 s) before and after Aβ42 application in Control BEC (Before: 5 mice, 8 arterioles, 24 stimulations, After: 5 mice, 8 arterioles, 32 stimulations) and CD2AP BEC-KD (Before: 6 mice, 15 arterioles, 37 stimulations, After: 6 mice, 15 arterioles, 43 stimulations) mice. G) Maximum dilation of penetrating arterioles after whisker stimulation Control BEC and CD2AP BEC-KD mice. Each dot represents one stimulation. *One-way ANOVA followed by Tukey Post Hoc analysis. H) Overall response of penetrating arterioles to whisker stimulation Control BEC and CD2AP BEC-KD mice. Each dot represents one stimulation. *Welch’s ANOVA followed by Dunnett’s T3 Post Hoc analysis. Power spectral density of arteriole from I) Control BEC and J) CD2AP BEC-KD mice before (Control: 5 mice, 11 arterioles, CD2AP BEC-KD: 6 mice, 15 arterioles) and after (Control: 5 mice, 13 arterioles, CD2AP BEC-KD: 6 mice, 15 arterioles) the application of Aβ. K) Maximum peak height around 0.1 Hz (0.075-0.125 Hz) from Control BEC and CD2AP BEC-KD mice before and after the application of Aβ. * Each dot represent one arteriole, *Two-way ANOVA. ^#^Unpaired Student t-test with Welch correction. L) Spectrograms of arteriole vasomotion from Control BEC and CD2AP BEC-KD mice before and after the application of Aβ. M) Capillary red blood cells speed before (n = 5 mice, 14 recordings) and after Aβ application (5 mice, 9 recordings) in Control BEC mice and before (6 mice, 15 recordings) and after Aβ application (6 mice, 15 recordings) in CD2AP BEC-KD mice. Each dot represents one recording. *Two-way ANOVA. ^#^Unpaired Student t-test with Welch correction. N) Representative example of the capillary line scan in Control BEC and CD2AP BEC-KD mice before and after the Aβ application. Data are presented as mean ± SEM. *P<0.05, **P<0.001, ***P<0.001, ****P<0.0001, ^#^P<0.05.

Aβ is a powerful vasoactive agent with multiple cellular targets including mural cells (Nortley et al., 2019), astrocytes (Sanchez-Mico et al., 2021), neurons (Mucke and Selkoe, 2012) and vascular endothelial cells (Nikolakopoulou et al., 2021). We therefore tested the effects of Aβ42 oligomers on brain vessels of CD2AP BEC-KD and control animals. visualized under awake two-photon microscopy for 20 min (Figure 5E). Previous studies have shown that Aβ can constrict brain vessels (Iadecola, 2013; Nortley et al., 2019). Similarly, we observed a constriction of penetrating arteriole in response to Aβ. However, no difference was observed between CD2AP BEC-KD and controls (Figure S9). Next, we examined the impact of CD2AP knockdown in BECs on functional hyperemia before and after Aβ superfusion (Figure 5F). In line with our previous observations (Figure 2 and 3), we confirmed the reduced overall response to whisker stimulation in CD2AP BEC-KD mice compared to control animals (Figure 5H). We also observed reduced vasodilation upon whisker stimulation in control animals in the presence of Aβ42 oligomers (Figure 5G and H), consistent with a report showing that Aβ40 impairs neurovascular coupling (Iadecola, 2013). The magnitude of this effect was similar to that of CD2AP BEC-KD mice. Importantly, we found that Aβ42 oligomers no longer influenced functional hyperemia when applied on the brain surface of CD2AP BEC-KD mice (Figure 5G and H). This result indicates that the lack of endothelial CD2AP occludes the response to challenges such as Aβ.

Next, we evaluated vasomotion (Figure 5I to L) – rhythmic oscillations in arteriole diameter at ∼0.1Hz – a process that could contribute to the paravascular clearance of Aβ through the glymphatic/lymphatic system and is altered in mouse model of amyloidosis (van Veluw et al., 2020). The maximum peak height during vasomotion was affected when all the groups were compared according to genotype and Aβ application (Figure 5K). Specifically, Aβ alone had no effect on vasomotion in control BEC mice. However, Aβ significantly reduced vasomotion only when CD2AP was depleted (Figure 5K). Finally, we evaluated the effect of Aβ on the capillary resting red blood cell speed on the basis that Aβ42 oligomers were previously shown to constrict capillaries, a process that could contribute to reduced CBF in AD (Korte et al., 2020). We found a reduction in red blood cells speed in CD2AP BEC-KD mice when compared to that of control animals (Figure 5M and N). This finding aligns with the lower CBF that we observed in CD2AP BEC-KD mice (Figure 2). When comparing the groups individually, Aβ42 oligomers had no impact on the speed of capillary red blood cells in control mice but caused a reduction in speed in CD2AP BEC-KD animals (Figure 5M and N). Together these data highlight that the loss of endothelial CD2AP increases Aβ40 accumulation, reduces vasomotion and increases the sensitivity of specific vessels to Aβ for decreasing resting CBF and vasomotion.

### Reelin prevents A**β**-mediated cerebrovascular dysfunction in capillaries and arterioles of endothelial CD2AP-depleted animals

Reelin protects against Aβ toxicity in neuronal cells (Durakoglugil et al., 2009). Therefore, we asked the question whether Reelin can prevent cerebrovascular dysfunction caused by Aβ in the presence or absence of brain endothelial CD2AP. Specifically, we evaluated functional hyperemia, vasomotion and resting capillary red blood cells speed before and during the superfusion of Reelin alone or in combination with Aβ in control and CD2AP BEC-KD mice. Reelin was superfused for 20 min prior to the co-superfusion of Aβ + Reelin during two-photon imaging (Figure 6A). No difference was found between control and CD2AP BEC-KD mice for the response of arterioles to Reelin or Reelin + Aβ (Figure S10). When comparing all groups, we observed an effect of genotype on the overall response during functional hyperemia (Figure 6B to D). Specifically in control mice, Reelin did not prevent the negative impact of Aβ on functional hyperemia in penetrating arterioles: the overall response to whisker stimulation was lower after the superfusion of Reelin and Aβ than before the drugs (Figure 6B and D). Consistent with our finding that Aβ alone has no effect on neurovascular coupling in the CD2AP BEC-KD mice (Figure 5), co-treatment of Aβ and Reelin did not elicit any effect in the same group of mice (Figure 6D). When comparing all groups, we observed an effect of genotype on vasomotion (Figure 6G). After Reelin superfusion, vasomotion was lower in CD2AP BEC-KD mice than in controls (Figure 6G). Importantly, co-treatment of Aβ and Reelin in CD2AP BEC-KD mice resulted in increased vasomotion compared to Reelin alone (Figure 6G). This was not observed in controls animals, further supporting our finding that Reelin signaling is altered upon loss of endothelial CD2AP (Figure 3). Since Aβ reduces vasomotion in CD2AP BEC-KD animals, this result indicates that Reelin prevented the reduction in vasomotion induced by Aβ when CD2AP is depleted in the endothelium. Finally, we confirmed the reduced capillary red blood cells speed in CD2AP BEC-KD animals compared to controls (Figure 6I and J). Further, Reelin increased resting CBF in CD2AP BEC-KD mice (Figure 6I and J), reinforcing the possible effect of this glycoprotein on vessel dilation (Figure 3, 4 and S6). Strikingly, in CD2AP BEC-KD mice, the capillary red blood cell speed remained higher than the baseline when Aβ was co-applied with Reelin, a phenomenon that was not observed in the controls (Figure 6I and J). The capillary red blood cells speed was not different between the CD2AP BEC-KD and control animals after Reelin or Reelin + Aβ treatment (Figure 6I and J). In summary, Reelin helps to prevent some of the noxious effects of Aβ on CBF and vasomotion in mice with lower brain endothelial CD2AP.

**Figure 6:**
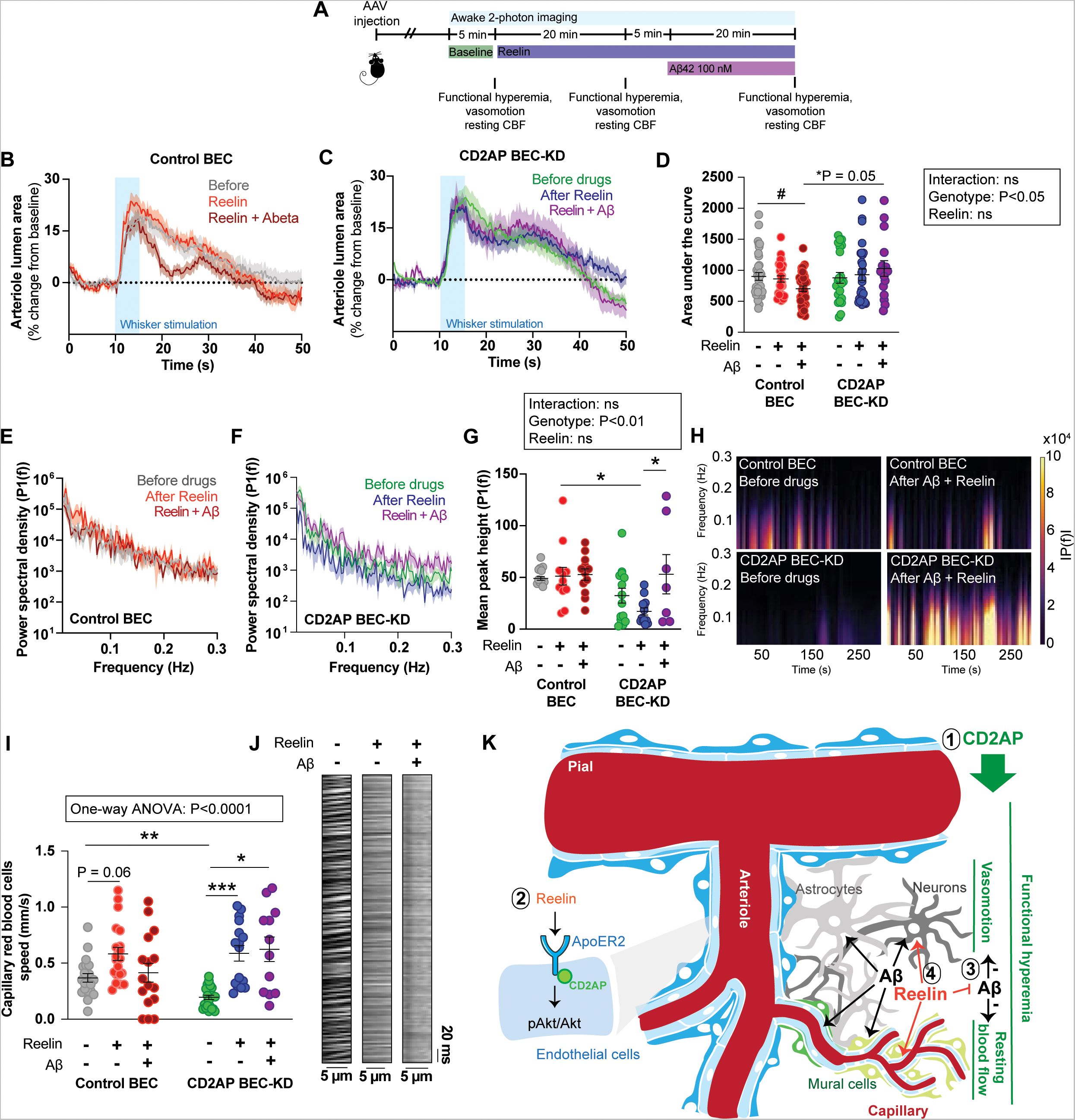
Reelin protects against the effect of A**β** on capillary resting blood flow and vasomotion in CD2AP BEC-KD mice. A) Timeline of the drug application during the *in vivo* experiments. Penetrating arterioles transversal luminal area in response to whisker stimulation (5 s) before and after Aβ42 application of B) Control BEC (Before: 6 mice; 13 arterioles, 35 stimulations, after Reelin: 6 mice; 13 arterioles, 27 stimulations, after Reelin + Aβ: 6 mice; 13 arterioles 37 stimulations) and C) CD2AP BEC-KD mice (Before: 7 mice, 11 arterioles, 25 stimulations, after Reelin: 7 mice, 12 arterioles, 30 stimulations, after Reelin + Aβ: 7 mice, 7 arterioles, 19 stimulations). D) Overall response of penetrating arterioles to whisker stimulation in Control BEC and CD2AP BEC-KD mice. Each dot represents one stimulation. *Two-way ANOVA followed by Sidak’s multiple comparisons. ^#^One-way ANOVA followed by Tukey Post Hoc analysis. E) Power spectral density of arteriole from Control BEC mice before (6 mice, 11 arterioles) and after the application of Reelin (6 mice, 12 arterioles) or Aβ + Reelin (6 mice, 12 arterioles). F) Power spectral density of arteriole from CD2AP BEC-KD mice before and after the application of Reelin or Aβ + Reelin (Before Reelin: 7 mice, 14 arterioles, After Reelin: 7 mice, 13 arterioles, After Aβ + Reelin: 5 mice, 7 arterioles). G) Mean peak height around 0.1 Hz (0.075-0.125 Hz) from Control BEC and CD2AP BEC-KD mice before and after the application of Reelin or Aβ + Reelin. Each dot represents one vessel. *Two-way ANOVA. H) Representative spectrograms from Control BEC and CD2AP BEC-KD mice before and after the application of Aβ + Reelin. I) Capillary red blood cells speed in Control BEC and CD2AP BEC-KD mice before (Control BEC: 6 mice, 20 recordings; CD2AP BEC-KD: 7 mice, 23 recordings), and after Reelin (Control BEC: 6 mice, 20 recordings; CD2AP BEC-KD: 4 mice, 16 recordings) or Reelin + Aβ (Control BEC: 5 mice, 16 recordings; CD2AP BEC-KD: 4 mice, 12 recordings). application. Each dot represents one recording. *Welch’s ANOVA followed with a Dunnett’s T3 Post Hoc analysis. J) Representative example of the capillary line scan in CD2AP BEC-KD mice before and after the Reelin and Reelin + Aβ application. K) Summary of the findings of the study. 1. Endothelial CD2AP is involved in functional hyperemia, vasomotion and resting blood flow. 2. In brain endothelial cells, CD2AP regulates ApoER2 levels and Reelin signaling. 3) Loss of CD2AP in the endothelium worsen the effect of Aβ on resting blood flow and vasomotion. 4) Reelin prevent the effect of Aβ on resting blood flow and vasomotion. Data are presented as mean ± SEM. *P<0.05, **P<0.01, ***P<0.001, ****P<0.0001, ^#^P<0.05

## Discussion

Mounting evidence highlights the importance of cerebrovascular defects in the pathogenesis of AD: Deregulation of vascular factors appears in the earliest stage of disease making the brain more vulnerable to insults such as Aβ (Zlokovic, 2005). Here, we studied the role of CD2AP in the brain vasculature in the context of AD. Our results indicate that 1) endothelial CD2AP is important for cerebrovascular and memory functions; 2) the endothelial form regulates Reelin-ApoER2 signaling *in vitro* and *in vivo*; 3) loss of CD2AP in the brain endothelium renders vessels more vulnerable to Aβ; and 4) the toxicity induced by Aβ on cerebrovascular functions can be partially alleviated with Reelin.

BECs respond directly to neuronal activation and release molecules that control vessel tone and blood flow (Chen et al., 2014; Chow et al., 2020; Longden et al., 2021). Here we showed that loss of endothelial CD2AP alters neurovascular coupling, resting blood flow and vasomotion at basal level, and changes the response to Aβ. These defects reflect the vast and complex vascular dysfunction seen in AD (Schaeffer and Iadecola, 2021). While CD2AP depletion reduces ApoER2 expression in BECs, Reelin-ApoER2 signaling monitored with Akt activation was enhanced. Further. Reelin increased vessel dilation and resting CBF in animals with lower BEC CD2AP. The causes underlying the defects in CD2AP-depleted BECs remain to be elucidated but they hint to other mechanisms controlled by CD2AP. For instance, in addition to Dab1, ApoER2 can also associate with Dab2 that is one of the strongest binding partners for CD2AP (Rouka et al., 2015). Signaling through Dab2 adaptor activates Akt and decreases eNOS activation (Sacharidou et al., 2018), a potent vasodilator. Thus, loss of CD2AP might increase ApoER2/Dab2-mediated Reelin signaling in the endothelium. Alternatively, in a non-mutually exclusive scenario, based on its cytoskeletal and membrane trafficking roles, CD2AP may be important for the formation and anchorage of caveolea (Chow et al., 2020; Longden et al., 2021) that are critical for blood flow regulation. While we uncover a functional Reelin-ApoER2-CD2AP axis in the brain endothelium and that endothelial CD2AP is critical for modulation of cerebrovascular function, we can also not exclude the possibility that Reelin superfusion and/or Aβ treatment affect(s) directly neuronal activity with repercussions on the vascular system. In any case, we demonstrate that the presence of CD2AP in the endothelium is key in regulating the interface between the vascular and neuronal systems under physiological and pathological conditions.

The present results suggest that CD2AP achieves vascular protection in AD. Data gathered in postmortem human samples and in mice indicate that downregulation of CD2AP may be involved in cognitive symptoms in AD, but animal data further shows that CD2AP is a key player in brain vascular processes (CBF, vasomotion and neurovascular coupling). Taking our findings in the context of the two-hit model (Zlokovic, 2005), the loss of CD2AP in BECs (1^st^ hit) would weaken brain vascular function and predispose the brain vasculature and parenchyma to a second hit mediated by Aβ and tau. Our findings of increased sensitivity of brain vessels lacking CD2AP to Aβ and of increased formation of Aβ plaques in CD2AP EC-KO, are in support of this working model. However, since the protein is expressed ubiquitously, the role of CD2AP in AD is definitely more complex: CD2AP is involved in the processing of amyloid precursor protein in neurons (Ubelmann et al., 2017) and may have uncovered functions in glial cells. Therefore, loss of vascular CD2AP might drive the initial stage of the disease but its association with plaques accumulation and insoluble tau in the later stages would imply its deregulation in other cell types.

In humans, genetic changes in *CD2AP* are associated with both focal segmental glomerulosclerosis, a leading cause of proteinuric kidney disease that progresses to end-stage renal failure (Kim et al., 2003) and to AD (Bertram et al., 2007). The brain and the kidney are both densely vascularized where organs are very vulnerable to small vessel dysfunction (Sweeney et al., 2019; Viggiano et al., 2020) and diversity of the vasculature is a key element in tissue homeostasis. In our CD2AP EC-KO, the kidney appears to be spared although we cannot exclude that subtle changes in the tissue could contribute to the phenotypes observed in the brain. It is known that the two main causes of chronic kidney disease are diabetes and high blood pressure (Chen et al., 2019), and both are key risk factors for AD (Kivipelto et al., 2018). Further, chronic kidney disease also predicts cognitive decline (Buchman et al., 2009). Given the role of CD2AP on vascular health unveiled here, its dysfunction could compromise either or both organs, depending on the genetic and environmental factors involved. Accordingly, genes linked to the kidney system are emerging as risk factors for AD (Kunkle et al., 2020) while patients with chronic kidney disease are predisposed to cognitive dysfunction and dementia (Elias et al., 2013; Viggiano et al., 2020; Zammit et al., 2016).

In summary, the present work unveils a hitherto unknown protective role of CD2AP against cerebrovascular dysfunction by maintaining the essential processes of CBF in distinct vascular segments. Developing therapeutic strategies to boost Reelin-CD2AP-ApoER2 pathway in cells within the brain vasculature may offer a refined therapeutic avenue for the treatment of AD.

## Methods

### Human brain samples

The brain parietal cortices samples used for this study were provided from the cohort of volunteers from the Religious Order Study, a longitudinal clinical-pathologic study of aging and dementia. An extensive amount of clinical and neuropathological data published from that cohort are available (Bennett, 2006; Tremblay et al., 2007; Tremblay et al., 2017). Volunteers characteristics are available in Table S1 and were published previously (Tremblay et al., 2011; Tremblay et al., 2017). Participants to this study were recruited without known dementia and underwent a clinical evaluation every year until death. Cognitive function of participants was evaluated using 21 cognitive performance test that were evaluated by a clinical neuropsychologist and expert clinician (Bennett et al., 2006). AD diagnosis was assessed based on a significant decline in at least two cognitive domains including episodic memory. Participants were classified in the MCI group if they showed cognitive impairment but were not diagnosed with dementia (Bennett et al., 2002) and NCI volunteers had no cognitive impairment (Bennett et al., 2012). Using 19 cognitive tests, a global cognitive score and a score for five different cognitive domains (episodic, semantic and working memory, perceptual speed and visual-spatial ability) was generated (Wilson et al., 2002). The volunteers were interviewed about use of anti-hypertensive and diabetes medication during the two weeks before each evaluation (Arvanitakis et al., 2008; Arvanitakis et al., 2016). The neuropathological diagnosis was based on a blinded evaluation of the ABC scoring method from the National Institute of Aging – Alzheimer’s Association (NIA-AA) guideline (Montine et al., 2012): A) Thal score (accumulation of Aβ plaques) (Thal et al., 2002), B) Braak score (neurofibrillary tangle pathology) (Braak and Braak, 1991), and C) CERAD score (neuritic plaques pathology) (Mirra et al., 1991).

#### Preparation of whole homogenates and microvessel-enriched extracts from human parietal cortex

Cortical extracts were prepared as described previously (Tremblay et al., 2011). Brain microvessel-enriched extracts from frozen human parietal cortex samples were prepared according to Bourassa et al., (Bourassa et al., 2019; Bourassa et al., 2020). Briefly, parietal cortex samples were thawed on ice in a microvessel isolation buffer (MIB; 15 mM HEPES, 147 mM NaCl, 4 mM KCl, 3 mM CaCl_2_ and 12 mM MgCl_2_) containing protease and phosphatase inhibitor cocktails (Bimake, Houston, TX), meninges and white matter were removed, and were then homogenized in MIB and spun at 1,000 g for 10 minutes at 4°C. The pellet was then homogenized in MIB containing 18% dextran (from leuconostoc mesenteroides, M.W. 60,000 – 90,000; Sigma-Aldrich) and centrifuged at 4,000 g for 20 min. at 4°C. The pellet was resuspended in MIB and filtered through a 20 µm nylon filter (Millipore). Vessels, retained on the filter, were deposited on glass slides for immunofluorescence staining or homogenized in lysis buffer (150 mM NaCl, 10 mM NaH_2_PO_4_, 1% Triton X-100, 0.5% SDS and 0.5% sodium deoxycholate), sonicated and spun at 100,000 g for 20 min. at 4°C. The resulting supernatant was concentrated using a Vivaspin device (MWCO, 3 kDa; Sartorius Stedim Biotech) and kept for Western immunoblotting analyses. In addition, the eluate containing the post-vascular fraction was pelleted following centrifugation at 16,000 g for 20 min. at 4°C, homogenized in lysis buffer and spun again at 100,000 g for 20 min. at 4°C. The resulting supernatant was preserved for Western immunoblotting analyses. Protein concentrations in all fractions were measured using the bicinchoninic acid assay (Thermo Fisher Scientific). 1 NCI, 2 MCI and 1 AD samples were lost during the sample preparation.

### Animals

Four different mouse lines were used for the current study: C57BL6 (77 males), PDGFRβ-Cre x RCL-GCaMP6s mice (10 females, 16 males), CD2AP.flox.cko.C-H05-A10-1_Tie2.Cre.tg (Control EC and CD2AP EC-KO, 26 males, 4 females) and CD2AP.flox.cko.C-H05-A10-1_Tie2.Cre.tg_Thy-1.PrP.hu.APP.hu.PS2.tg (PS2APP Control EC and PS2APP CD2AP EC-KO, 34 males).

CD2AP.flox.cko.C-H05-A10-1_Tie2.Cre.tg_Thy-1.PrP.hu.APP.hu.PS2.tg mice were created by crossing: Thy-1.PrP.hu.APP.hu.PS2.tg.B6 (C57BL/6J background), Map4K4.flox.cko.1B5-1_Tie2.Cre.tg (C57BL/6N background) and CD2AP.flox.cko.B6N.C-H05-A10-1 (C57BL/6J background). The PS2APP mice express the human PS2 gene with the N141l mutation and the human APP gene with the Swedish mutation (Meilandt et al., 2020). All mice were maintained in a room kept at 22°C with a 12h light cycle and had ad libitum access water and standard chow diet. Animal procedures for the C57BL6, Control EC and CD2AP EC-KO mice were performed in accordance with the Canadian Council on Animal Care guidelines and were approved by the Health Sciences Animal Care Committee at the University of Calgary. Animal care and handling procedures for PS2APP Control-EC and PS2APP CD2AP EC-KO were reviewed and approved by the Genentech Institutional Animal Care and Use Committee and were conducted in full compliance with regulatory statutes, Institutional Animal Care and Use Committee policies, and National Institutes of Health guidelines.

### AAV preparation

#### Plasmid construction

The DNA sequences encoding the mouse U6 promoter, CMV-eGFP and WPRE from pLL3.7 (a gift from Luk Parijs (Addgene plasmid # 11795; http://n2t.net/addgene:11795; RRID:Addgene_11795) (Rubinson et al., 2003) were subcloned into the MluI and XhoI restriction sites of pENN.AAV.hSyn.Cre.WPRE.hGH (a gift from James M. Wilson (Addgene plasmid # 105553; http://n2t.net/addgene:105553; RRID:Addgene_105553)), replacing the hSyn.cre.WPRE with U6.CMV.eGFP.WPRE to generate pAAV.U6.CMV.GFP.WPRE.hGH. The siRNA sequences purchased from Santa Cruz Biotechnology and Dharmacon that had previously been validated *in vitro* to efficiently knock down CD2ap, were engineered into shRNA 97-mer backbones (Pelossof et al., 2017), synthesized into a large primer and inserted immediately downstream of the U6 promoter to generate pAAV.U6.shRNA.CMV.GFP.WPRE.hGH using the NEBuilder hifi DNA assembly cloning kit (New England Biolabs). Corresponding constructs with tdTomato as the reporter were generated by swapping in the 5’-MluI and 3’ AgeI U6-shRNA-CMV fragment from each pAAV.U6.shRNA.CMV.GFP.WPRE.hGH construct in place of the corresponding sequence in pAAV-U6-shRNA-CAG-tdTomato (Batool et al., 2021) to generate pAAV.U6.shRNA.CMV.tdTomato.WPRE.hGH. The sequences of all constructs were verified by Sanger DNA sequencing.

#### AAV production

AAV viral vectors containing the BR.1 capsid (Körbelin et al., 2016) were generated using the methods of Challis et. al. (Challis et al., 2019). Briefly, 293FT cells (Thermofisher) were grown to ∼90% confluency in Corning hyperflasks (Corning) and co-transfected with 129 µg pHELPER (Agilent), 238 µg rep-cap plasmid encoding BR.1 (Körbelin et al., 2016) and 64.6 µg equimolar mixtures of pAAV.U6.shRNA.CMV.GFP.WPRE.hGH or pAAV.U6.shRNA.CMV.tdTomato.hGH corresponding to either the four Dharmacon shRNA sequences, the three Santa Cruz shRNA sequences or a scrambled control using the PEIpro transfection reagent (Polyplus). AAVs were precipitated from media harvested after 3 days and 5 days using 40%PEG/2.5M NaCl and pooled the cells harvested after 5 days in buffer containing 500 mM NaCl, 40 mM Tris Base and 10 mM MgCl_2_. The lysate was incubated with 100 U/mL salt-active nuclease (Arcticzymes) at 37°C for 1 h and then centrifuged at 2000 xg for 15 min. AAV was purified from the resulting lysate using an iodixanol step gradient containing 15, 25, 40 and 60% iodixanol (Batool et al., 2021) in optiseal tubes (Beckman) followed by centrifugation at 350,000 xg using a Type 70 Ti ultracentrifuge rotor (Beckman). Following centrifugation, the AAVs were harvested from the 40% layer using a 10 cc syringe and 16-guage needle, diluted in 1XPBS containing 0.001% pluronic F68 (Gibco) and filtered using a 0.2 um syringe filter. The AAVs were concentrated and buffer-exchanged by 5 rounds of centrifugation using Amicon Ultra-15 100 kDa molecular weight cut off centrifugal filter units (Millipore). The titer was determined using the qPCR Adeno-Associated Virus Titration kit (Applied Biological Materials) and the purity was verified by SDS-PAGE and total protein staining using instant blue reagent (Expedeon). The AAV was administered to 30 days old male C57BL6 mice or PDGFRβ-Cre x RCL-GCaMP6 mice with a retro-orbital injection (150 μl, 1X10^11^ GC).

### Behavioural tests

Mice were habituated by gentle handling for a minimum of two to three consecutive days prior to behavioural test. Animals were transported to the testing room to acclimatize for at least 30 minutes in their home cage prior to any test. Water maze was conducted with a minimum of 3 rest days before or after novel object recognition test, and we allotted a minimum of a week between the Barnes maze test (day 12) and any behavioural test.

#### Barnes Maze

The Barnes maze test was based on Sunyer et al., 2007 (Sunyer B, 2007). The Barnes circular maze task was conducted on a white circular table that is 1m in diameter, with 20 holes equally spaced around the perimeter. The test consisted of three stages: habituation, testing, and probe trials. During the habituation phase, mice were placed in the middle of the table and gently guided to the target hole containing the escape box. The mice were kept in the escape box for two minutes in the dark.

In the testing phase, mice were placed in the middle of the maze and allowed to explore the maze for three minutes. During this time, the number of non-escape hole approaches and latency to reach the escape box were recorded. The trial ended when three minutes have passed or when the mouse entered the escape box. If the mice did not reach the target hole in time, they were again guided to the box. Once the mice were inside the escape box, the light source was turned off and they were kept in the box for one additional minute. Animals underwent four consecutive days of trial, consisting of four trials per day, with an inter-trial interval of 15 minutes. After each trial, the apparatus was cleaned with 70% ethanol to eliminate any potential odor cues.

The probe trial was conducted on day 5 and again on day 12 with the target hole closed. Mice were allowed to explore the maze for 60 seconds and the number of head pokes into non-escape holes and latency to reach the target hole were recorded.

The Barnes Maze was done 12 weeks after the AAVBR1 injection in the Control BEC and CD2AP BEC-KD mice and at 14 months of age in the Control EC and CD2AP EC-KO mice.

#### Morris Water Maze

Morris water maze was conducted using a circular tank (113cm) filled with water (24.5 ± 0.5°C). A hidden platform was placed 1 cm below the surface of the water and three cups of milk powder were added to the water to make it opaque. The mice underwent training trials, followed by probe trials. During each training trial, the mice were released into the maze in one of the four quadrants following a randomized order of quadrants that was consistent for all mice (*e.g.,* NSEW for Day 1 AM and WNES for Day 1 PM) and allowed to explore for 60 seconds. The trial ended when the 60 seconds elapsed or when the mice located the platform. If the platform was not located within the time, the animal was gently guided to the platform and left there for 10 seconds. Six training sessions took place: four trials AM and four trials PM, separated by four hours between AM and PM sessions over three days. The latency to locate the platform and time spent in the target quadrant was recorded using the ANYmaze software. Two probe trials were performed on the fourth day. First, a sole probe trial was conducted by removing the platform from the pool and allowing the mice to explore for 60 seconds. Following this, a single visible platform trial was conducted by placing a bright-colored object on the platform to determine if the mice had any visuomotor or motivational impairments in reaching the platform. The Morris Water Maze was done in 8 months-old Control EC and CD2AP EC-KO mice.

#### Novel object recognition test

We used the protocol from Miedel et al., 2017 (Miedel et al., 2017). The novel object recognition test is comprised of three stages: habituation, training, and testing. During the habituation stage, the mice were placed into an empty open box (49cm x 49cm) and allowed to explore for five minutes. 24 hours later, in the training stage, the mice were placed in the middle of the box, containing two identical objects placed at opposite quadrants of the box. Mice were allowed free exploration for 10 minutes and then returned to their holding cages. The testing phase took place 30 minutes after training and the mice were returned to the box now containing one familiar object and one novel object. Mice were again allowed free exploration for 10 minutes. The time spent exploring each object was recorded using ANYmaze software. Exploration of the object was defined as the mouse’s nose pointing towards the object and being within 2-3 cm from the object. For the Control EC and CD2AP EC-KO mice, the novel object recognition task was done at 8 and 16 months of age. For the Control BEC and CD2AP BEC-KD mice, the object recognition task was done 13 weeks after the AAVBR1 injection.

#### Open-field

The open-field test was used to examine locomotor function and anxiety. A blue open box measuring 49cm by 49cm was used. Mice were placed in the middle of the box and allowed to explore for 10 minutes. During the test, the amount of time and distance travelled in the center and outer area of the maze was recorded using the ANYmaze software. The openfield was done 14 weeks after the virus injection in the Control BEC and the CD2AP BEC-KD and at 8 and 15 months of age in the Control EC and CD2AP EC-KO mice.

### MRI

Mice were subjected to MRI according to a previously published protocol (Hashem et al., 2020). One mouse was excluded from the analysis because of a tumor. Briefly, mice were spontaneously ventilated with a mixture of 2% isoflurane, 70% N_2_ and 30% O_2_. The mouse was set up in the MRI system on top of a heating pad to maintain body temperature at 37°C. A thermometer with a rectal probe allowed to follow body temperature. Breathing rate was monitored continuously. Imaging was performed using a 9.4T, 210 mm bore Magnex magnet, run by Bruker Avance II hardware controlled by Bruker ParaVision 5.1 software (Bruker Avance console, Bruker BiospinGmbH, Rheinstetten, Germany).

Perfusion was measured by acquiring axial slices around bregma -1.94 mm using a CASL-HASTE sequence: TR = 3000 ms, TE = 13.5 ms, FOV = 25.6 × 25.6 mm, matrix size = 128 × 128 pixels, slice thickness = 1 mm, 16 averages. Two control and two tagged images were collected per measurement. Following these, a T1 map was obtained using a RARE-VTR sequence: TE = 10 ms, TR = 100, 500, 1000, 3000, and 7500 ms.

Quantification of the perfusion images and T1 map were done using a custom Matlab script (Johnson, 2017), in which CBF was calculated on a voxel-by-voxel with the following equation (Buxton, 2005; Pekar et al., 1996):

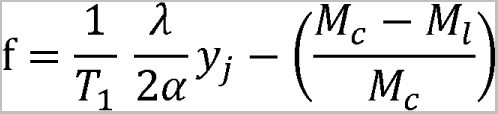

Were f is the tissue perfusion (ml/100 g/min.), λ(0.9) is the blood-brain partition coefficient (Herscovitch and Raichle, 1985; Leithner et al., 2010), *M_c_* is the average value for the control images, *M_l_* is the average value for the tagged images, α(0.675±0.044) is the efficiency of the spin labelling (Johnson, 2017) and *T*_1_ is the *T*_1_ value for each voxel. A ROI was generated usinga tagged image and was then used on the T_1_ map to obtain a mean ± SD for perfusion. Each ROI was traced 3 times and the values presented in the figure is the average of these 3 ROI. Control BEC and CD2AP BEC-KD mice were subjected to MRI 14 days and 21 days after AAV injection, Control EC and CD2AP EC-KO mice were imaged at 6 and 14 months of age. Resting cerebral blood flow values were similar to previous values published for the same brain regions (Morimoto et al., 2018; Munting et al., 2021).

### Reelin purification

Reelin was purified from conditioned media of HEK 293 cells stably transfected with Reelin cDNA (a gift from Dr. Joachim Herz laboratory, University of Texas Southwestern Medical Center, and Dr. Tom Curran, University of Pennsylvania) as per (Förster et al., 2002; Jiang et al., 2016; Kiroski et al., 2020; Weeber et al., 2002). Briefly, for the cell culture experiments, cells were cultured in low glucose DMEM (Gibco, Thermo Fisher, Waltham, M. 11885084) supplemented with FBS until they reached full confluency and then, the media was switched to phenol red-free Opti-MEM for 2 days. Afterward, the conditioned Opti-MEM was centrifuged at 5000g for 15 min to pellet any cell debris. The clarified media was then sterile filtered and concentrated by 30 times using Amicon Ultra-15 100 kDa Centrifugal Filter Units (EMD Millipore). For the pressure myography and the awake two-photon imaging experiments, after the cells reached full confluency, the media was switched to low glucose DMEM (Gibco, Thermo Fisher, Waltham, M. 11885084) supplemented with 0.01% bovine serum albumin (BSA) for 2 days. After, the media was centrifuged at 5000g for 15 min to pellet any cell debris. The supernantant was concentrated 30 times using Amicon Ultra-15 100 kDa Centrifugal Filter Units (EMD Millipore). The media was exchanged to buffer A (20 mM Tris, 2 mM CaCl_2_, 10 mM NaCl) and concentrated again 30 times. The concentrated media in buffer A was purified using G-Sepharose (Sigma) beads with buffer A from 0.1 M to 0.4 M NaCl concentration. The Reelin samples were collected between 0.25 M and 0.3 M NaCl. The samples were concentrated 30 times using Amicon Ultra-15 100 kDa Centrifugal Filter Units (EMD Millipore). The solution was diluted in PBS and concentrated again using an Amicon Ultra-4 50 kDa Centrifugal Filter Unit (EMD Millipore) to replace the media with PBS while retaining the dissolved Reelin. The control solution was prepared with the same procedure used for the Reelin solution but instead of using Reelin-transfected HEK 293 cells, non-transfected HEK 293 cells were used. The concentration of Reelin was measured by DC Assay (BioRad Laboratories), whereas presence of the glycoprotein in the purified fraction was confirmed on Western blot with anti-Reelin antibodies, and was verified with mass spectrometry according to published methods (Jiang et al., 2016; Kiroski et al., 2020). Reelin activity was confirmed by Western blot using antibodies directed against phospho-Serine 473 Akt, a downstream target of the Reelin pathway, in cell lines, as well as by comparing with that of a commercially available central active fragment of Reelin.

### A**β**42 oligomers solution

Human Aβ42 (HB9805, hellobio) was dissolved in PBS at 100 μM and incubated at 37°C for 72 hours (You et al., 2012). The solution was dissolved in aCSF at 1 μM ∼30 min before the two-photon microscopy experiment. Aβ was superfused on the brain of animals ∼10 to12 weeks after the AAVBR1 injection.

### Awake two-photon imaging

All procedure for awake two-photon imaging were performed according to a previously published protocol (Tran and Gordon, 2015; Tran et al., 2018). The Briefly, two-photon imaging was performed to evaluate the effect of drugs on vessel dilation, neurovascular coupling, vasomotion and capillary red blood cells speed. <u>Day 1, head bar installation</u>: Mice were kept under anesthesia with isoflurane and received a sub-cutaneous injection of buprenorphine (0.05 mg/kg) and enrofloxacin (2.5 mg/kg). Under aseptic conditions, the skin on top of the head was removed and the skull was cleaned with lactate ringer before a metal head bar (0.5 g) was glued on the occipital bone using fast glue. The head bar position was secured on the brain using first 2 layers of a three components dental glue (C&B Metabond, Parkell Inc, NY, USA) and dental cement (OrthoJet Acrylic Resin, Land Dental MFG. CO., Inc., IL, USA) was added to form a well large enough to receive the microscope objective. A cut 16G needle was inserted in the well to allow drug superfusion during the imaging session. <u>Day 2 and 3, training sessions</u>: After 24h of recovery, mice were habituated to run on an air-supported spherical Styrofoam ball. Mice were head-fixed and were allowed to run for 15 min after which, intermittent air stimulation was delivered to the contralateral whiskers (15 stimulations, 5 s, every 60 s). <u>Day 4, craniotomy</u>: Mice were anesthetized with isoflurane and received an injection of buprenorphine for pain control (0.05 mg/kg). The bone and the dura were removed gently over the barrel cortex with constant immersion under HEPES CSF (142 mM NaCl, 5 mM KCl, 10 mM Glucose, 10 mM HEPES, 3.1 mM CaCl_2_, 1.3 mM MgCl_2_). After the surgery was completed, the cranial window was sealed with a cover glass with nine 0.175 mm holes using fast glue. The well was constantly maintained under aCSF to prevent the brain surface from drying. For some of the experiments, animals received a 150 μl retro-orbital injection of 5% FITC-dextran (MW 2,000,000, Sigma) dissolved in lactate ringer. The animals were then transferred to the microscope and imaging was started about 45 min later.

#### Drug application during awake two-photon imaging

The brain surface was superfused with aCSF at 0.9 ml/min with a carbogen pressurized system (95% O_2_, 5% CO_2_). The drugs were pre-dissolved in aCSF and setup on a syringe pump adjusted at 0.1 ml/min, allowing a 1 in 10 dilution of the drug applied on the brain surface.

#### Functional hyperemia

Pial arteriole were first distinguished from venules by vasomotion and response to whisker stimulations. The response of either pial vessels, connected penetrating arterioles (∼ 45 to 110 μm depth) or 4rth order capillaries (∼ 30 to 250 μm depth) to contralateral whisker stimulation was recorded. An air puff (5 s) was delivered through two glass tubes to the contralateral whiskers after recording a 10 s baseline. Response to whisker stimulation was recorded three times before and after each drug was applied.

#### Two-Photon Fluorescence Microscopy

Imaging was performed using a custom built two-photon microscope (Rosenegger et al., 2014) fed by a Ti:Sapph laser source (Coherent Ultra II, ∼4 W average output at 800 nm, ∼80 MHz). Image data were acquired using MatLab (2013) running an open source scanning microscope control software called ScanImage (version 3.81, HHMI/Janelia Farms)(Pologruto et al., 2003). Imaging was performed at an excitation wavelength of 800 nm for FITC-dextran or 900 nm for experiments involving GFP. The microscope was equipped with a primary dichroic mirror at 695 nm and green and red fluorescence was split and filtered using a secondary dichroic at 560 nm and two bandpass emission filters: 525-40 nm and 605-70 nm (Chroma Technologies).

#### Images analysis

##### FITC-injected animals

Images were analyzed using ImageJ. Vessel area was obtained from the FITC-labelled lumen. Each image was thresholded and a particle analysis was performed. No animals were excluded from the analysis.

##### AAV-injected mice

A fluorescent dye (Rhodamine-B or Texas red dextran) was i.v. injected to improve visualization of the vasculature and to allow us to measure pial or penetrator diameter changes to whisker stimulation, or red blood cell movement through fourth order capillaries.

*Capillary red blood cells speed analysis*.

#### Training a U-Net for RBC Flow Classification

A convolutional neural network, with the classical U-Net topology (Ronneberger) was trained to perform an automated segmentation to identify putative red blood cells. Briefly, a U-net acts somewhat similarly to a neural autoencoder to perform segmentation. The input data is bottlenecked through a narrow layer, forcing the convolutional neural network to identify latent structures/patterns that repeat in the data. The U-Net was pre-configured with the MATLAB (2021a) *unetLayers* function. The networks were trained and run with the Machine Learning toolbox in MATLAB 2021a. Given the large differences between flow rates in the Control BEC and CD2AP-BEC-KD condition, we trained two separate U-Nets for each data type. We refer to these as Unet-Slow and Unet-Fast. The output of the unit is a binary matrix (*M_k_*) of the original size as the input image *I_k_*. The matrix entry, *M_k_*(*i,j*) = 1 if pixel *i, j* in image *k* is part of a red-blood cell, and 0 otherwise. After the U-Net segmentation, distinct (isolated) cells were identified with the MATLAB function *bwconcomp.* This function assigns distinct integers to every individual RBC identified. For U-Net-slow the initial learning rate was, ε = 10^-4^ with an epoch size of 2346 iterations. RMSprop was implemented in a batch mode, with a batch size of 2 images due to RAM limitations. For Unet-slow, 2 epochs of training were used with the validation accuracy being 92.33% and the training accuracy being 94.09% at the conclusion of training. For U-Net-Fast, 30 epochs of training were used with a smaller number of iterations per epoch (84) owing to the smaller labeled data set. Further, the learning rate was halved after 20 epochs. The training error rate for U-Net-Fast was 86.55%. Due to the small training sample size, the data was not segregated into a validation set. Human validation to confirm network performance was utilized after learning. The training was robust to a variety of different conditions including smaller iterations per epoch, faster learning rate reductions, etc. The performance of both U-Nets on a sample of training frames is shown in Figures S10 and S11.

#### Supervisor Construction and Data Augmentation

To facilitate network generalization, we performed data-augmentation on the initially limited, hand-labeled data sets. Every labeled training image and its corresponding mask supervisor was flipped vertically, flipped horizontally, transposed, with the transposition also subsequently applied to the flips. Then, a GUI was created allowing a user to manually identify the number of cells and draw a mask around each individual cell. The mask was generated with the *inpolygon* function to set all the indices (*i,j*) within the drawn mask to *M_k_*(*i,j*) = 1. The mask matrix *M_k_*(*i,j*) was used as the supervisor for each individual frame *l_k_*(*i,j*). A total of 4692 images were used to train U-net slow (inclusive of data-augmentation). These correspond to 1173 images that were manually labeled. A total of 168 training images were used for U-Net Fast. A total of 129 images were used as validation data for U-Net slow during training.

#### Manual Correction of Segmented Data

While each U-Net was accurate in classifying RBCs, we created a graphical-user-interface that would plot *n* x *n* frames simultaneously, along with the segmentation mask overlayed on the original image. The user could then curate poorly classified cells out of the pool by clicking on them. This was added to avoid edge artefacts or poorly resolved parts of frames.

#### Estimation of Δx and Δt

For each image stack, the user would manually specify two vertical lines (Δx?) that would form an intersection point for the segmentation mask. This corresponds to a fixed index in the image matrix matrix *I_k_*(*i, j*_1_) and *I_k_*(*i, j*_2_). This allows for the simultaneous removal of edge effects in the u-net segmentation, and systematic error occurring due to edge occlusion artefacts. Then, Δx and Δt were estimated by

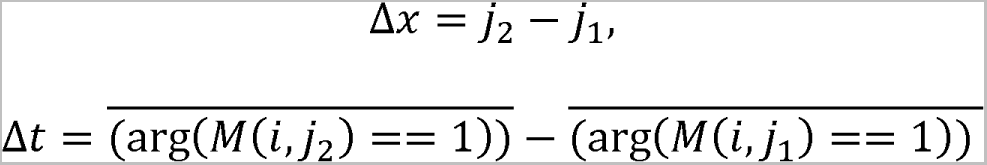

where the overbar denotes the finite average. Verbally, all pixels with a U-Net classification of 1 and an index of *j_2_* (*j*_1_) had their x-coordinate index (*i's*) averaged. The *Δt* is the difference between these two averages. An image of the final-output of this processing pipeline is shown below in Supplementary Figure S11 and S12. Data generated from 30 manual analysis using ImageJ had a 0.05% difference compared to the U-Net ouput from the same images. *Δx/Δt* was used to determine the capillary red blood cells speed (Kleinfeld et al., 1998).

#### Blood Vessel Tracking via REGDEMONs

The RBC/Astrocytes were tracked with a custom written code based on the Diffeomorphic Demon algorithm from (Vercauteren et al., 2009). More details on the exact implementation can be found in our previous work (Haidey et al., 2021).

#### Vasomotion analysis

We computed discrete Fourier transforms on a trace-by-trace basis for each recording and computed the power spectral density (PSD) in a vasomotion-related range, 0.01-0.3Hz. Raw traces were filtered in this range and a custom MATLAB script was prepared to extract mean and mode peaks from the filtered traces. To visualize spectral changes over time, we also computed Hamming-windowed short-time Fourier transforms for 20-second segments of each trace with 90% overlap between segments. Individual PSD profiles and spectrograms from each recording were then averaged to allow comparisons between groups.

#### Calculations

For the responses to drug superfusion and whisker stimulation during awake two-photon imaging, the vessel area was first analyzed as described in the ‘image analysis’ section. The average area was calculated for the baseline (5 sec for the whisker stimulation and 5 min for the drug superfusion). The area value after the baseline were subtracted from the baseline and a ratio was made on the baseline value and multiplied by 100: Vessel size (%) = area baseline – (area baseline – area) / area baseline x 100. The maximum dilation represents the peak area for individual experiments after the baseline. For the drug application and vasomotion, each value represents one vessel. For the whisker stimulation, the stimulations were done 3 times for each vessel. Data are presented as number of stimulations. The number of vessels and animals used are specified in the figure legend for each experiment.

#### Representative images for figures

Every image presented in the figures for the drugs application and the response to whisker stimulation were processed the same way. Two images were averaged and a Gaussian filter (0.5 or 1 pixel radius) was performed by ImageJ plugins.

### Pressure Myography

#### Vessel Isolation

Male C57BL/6 mice between 6 and 8 weeks of age were injected intraperioneally with sodium pentobarbital (50 mg/kg) to induce surgical anesthesia (i.e., stage 3, loss of blink reflex). A total of 12 animals were used to obtain the cerebral arteries (posterior and middle cerebral) described in this study. No animals were excluded from the analysis. To isolate cerebral arteries, anesthetized animals were first euthanized by decapitation, and the excised brain was placed in a cooled dissection chamber.

#### Arterial Pressure Myography

After isolation, arteries were cannulated on glass pipettes fitted in a pressure myography chamber (Living Systems, Burlington, VT). The vessel lumen was filled with Kreb’s buffer (115 NaCl, 5 mM KCl, 25 mM NaHCO_3_, 1.2 mM MgCl_2_, 2.5 mM CaCl_2_, 1.2 mM KH_2_PO_4_, and 10 mM D-glucose) containing 1% bovine serum albumin, pH was adjusted to 7.4 (Mishra et al., 2015; Mishra et al., 2018). The cannulated vessel/chamber apparatus was placed on the stage of an inverted microscope, and the vessel was superfused with Kreb’s buffer at a constant flow of 6-7ml/min using a peristaltic pump and suction line. Bath solution was maintained at 37^°^C and gassed with 95% air/5% carbon dioxide. After 5–10 minutes of equilibration, the intraluminal pressure of cannulated vessels was increased in a stepwise manner and then maintained at 70 mmHg. The vessels typically developed myogenic tone within 20–30 minutes. Continuous video measurement of the intraluminal vessel diameter was carried out using a diameter tracking system (IonOptix, Milton, MA). Drug-containing solutions were added to the bath through the perfusion pump. For endothelial denudation, an air bubble was slowly passed through the lumen of the vessel after cannulation of one end of the vessel. Physiological saline was then gently passed through the lumen to flush out any cellular debris. The vessel was then fully cannulated and allowed to equilibrate as described above before the application of intraluminal pressure and the development of myogenic tone. Endothelium loss was confirmed by loss of SKA-31 and bradykinin-evoked dilation in endothelium-stripped arteries; Figure S5 (Mishra et al., 2018). Drug induced-changes in vessel internal diameter were calculated and are presented as the absolute change (in microns) from the baseline intraluminal diameter recorded at steady state myogenic tone immediately prior to drug application.

#### Reagents

Bradykinin, EGTA (ethylene glycol-bis(2-aminoethylether)-N,N,N’,N’-tetraacetic acid), Phenylephrine hydrochloride ((R)-(-)-1-(3-hydroxyphenyl)-2-methylaminoethanol hydrochloride), (-)Norepinephrine bitartrate salt, 4-AP (4-aminopyridine), SNP (sodium nitroprusside), DMSO (dimethyl sulfoxide), Pinacidil (N-cyano-N0-4pyridinyl-N″-(1,2,2-trimethylpropyl)guanidine monohydrate), and all required chemicals were purchased from SigmaAldrich (Oakville, ON, Canada). Euthanyl (sodium pentobarbital, 250 mg/ml) was purchased from Bimeda-MTC Animal Health Inc, Cambridge, ON, Canada. SKA-31 (naphtho [1, 2-d] thiazole-2-ylamine) was synthesized as previously described (Sankaranarayanan et al., 2009). Reelin and MOCK media were prepared as described previously. SKA-31 and pinacidil were prepared as 10 mmol/L stock solutions in DMSO and then diluted directly into the external bath solution. The final concentration of DMSO reaching the tissue was typically 0.02% (vol/vol) or less. In preliminary experiments, we observed that a considerably higher concentration of DMSO (0.2% v/v final) had no effect on either basal myogenic tone or the responsiveness of cerebral arteries to vasodilator agents (Mishra et al., 2015).

### Cell cultures

Mouse primary BECs (mBECs) were prepared using an adapted protocol (Bourasset et al., 2009). No samples were excluded from the analysis. The brains of five C57BL6 mice aged between 2 and 6 months were used to cover a 75 cm^2^ flask. Mice were sacrificed under anesthesia with 2-bromo-2-chloro-1,1,1-trifluoro-ethane (Sigma) by cervical dislocation. The brains were placed in petri dish filled with ice-cold PBS where the cerebellum was removed and discarded. Meninges were removed by rolling brains on UV sterilized 30x30 cm kimwipes until they were no longer visible. The brains were transferred to a dry petri dish and diced thoroughly with a scalpel forming pieces about 1 mm^3^ in size. The homogenate was then added to the enzymatic digestion buffer (Worthington Biochemical Corp., 250 U/ml collagenase type II (LS004174), 10 U/ml dispase (LS02100) and 10 U/ml DNAase (LS006343) in DMEM) and incubated with agitation for 90 min at 37°C. After the digestion, the homogenate was pelleted by centrifugation at 200g for 3 min., the supernatant was aspirated and the pellet was resuspended in 4°C DMEM and pelleted again at 200g for 3 min. The supernatant was aspirated and brain material was homogenized in DMEM 20% BSA. The suspension was then centrifuged at 1500g for 20 min at 4°C pelleting the brain capillaries. The supernatant and floating white matter were aspirated and capillaries were resuspended in Medium 131 + MVGS supplement kit (Gibco, ThermoFisher, M-131-500 and S00525) and passed through a 70 μm cell strainer to dissociate the capillary cells. The capillary cell suspension was then supplemented with 4 μg/ml of puromycin (Sigma Aldrich, P8833-10MG) plated on collagen IV (Santa Cruz Biotechnology, sc-29010) coated culture plates and placed in a CO_2_ incubator at 37°C. After 2 days the media was changed to puromycin-free growth media and cells were cultured until reaching appropriate confluency.

hBECs (hCMEC/D3, a generous gift from Dr Amir Nezhad, University of Calgary) were cultured according to the supplier protocol (EMD Millipore, SCC066), in EndoGRO™-MV Complete Media Kit (EMD Millipore, SCME004). Cells were plated on Collagen I (EMD Millipore, 08-115) coated T75 culture flasks and passaged at 90% confluency.

#### Transfection of siRNA

hBECs and mBECs were transfected with human CD2AP siRNA (Santa Cruz Biotechnology, sc-29984) and mouse CD2AP siRNA (Santa Cruz Biotechnology, sc-29985), respectively. Both siRNA efficiently reduced the levels of CD2AP and its target, the ApoER2. A third efficient siRNA (ON-TARGETplus, Dharmacon, L-041647-01-0005) confirmed the knockdown of CD2AP in mBECs. For the mBECs, the transfection was performed 5 days after plating the cells, yielding a confluency of about 60%, using HiPerFect (QIAGEN) reagent according to the manufacturer instructions. For one well of a 24-well plate, 10 nM siRNA were added to the well. For both cell models, transfection media was aspirated and replaced with growth media after 3 hours. Cells were harvested, fixed or used for assays 42 hours and 7 days post-transfection for hBECs and mBECs, respectively.

#### Reelin treatment of hBECs

hBECs cells were first transfected with control or CD2AP siRNA according to the method describe above. After the transfection, hBECs cells were incubated for 60 min in DMEM media without FBS. The media was then replaced with DMEM with Reelin (20 nM) or control solution for 30 or 120 min.

### Immunostaining

#### Human brain samples

Microvessels from enriched fraction were dried on Superfrost Plus slides and fixed using a 10% formaldehyde solution (Sigma-Aldrich) 10 min at RT and then blocked with a 10% normal horse serum and 0.1% Triton X-100 solution in PBS for 1 h at RT. Then, samples were stained with goat anti-type IV collagen (Millipore, AB769, 1:500) and with rabbit anti-CD2AP (Sigma-Aldrich, HPA00326, 6μg/mL). Microvessels were then incubated with secondary antibodies (donkey anti-rabbit Alexa Fluor 555 or donkey anti-rabbit Alexa Fluor 647). Finally, cell nuclei were counterstained with DAPI and slides were mounted with Prolong Diamond mounting medium.

Between each step, three washes of 5 min with PBS 1X were performed. Images were taken using a laser scanning confocal microscope (Olympus IX81, FV1000; Ontario, Canada) and were acquired by sequential scanning using optimal z-separation at a magnification of 40X.

#### Cell culture, Control BEC-KD, CD2AP BEC-KD, Control EC and CD2AP EC-KO mice

Immunochemistry was performed based on published protocol (Jiang et al., 2016; Kiroski et al., 2020). Briefly, Control BEC-KD and CD2AP BEC-KD mice were anesthetized with ketamine/xylazine and perfused with ice-cold PBS followed 4% PFA. The brains were then rapidly dissected and post-fixed in 4% PFA for 24h, incubated in 30% sucrose for 48h. Control EC and CD2AP EC-KO mice were anesthetized with ketamine/xylazine and perfused with ice-cold PBS. The brains were then rapidly dissected and post-fixed in 4% PFA for 48h. After the post-fixation, brains from all mice were incubated in 30% sucrose for 48h, embedded in optimal cutting temperature (OCT) compound and frozen at -80°C until 20 μM thick coronal slices were performed for immunostaining. Sections were washed with PBS, blocked for 1h (1% goat serum, 0.1% Triton X-100) and incubated in primary antibody overnight. The next day, sections were incubated in the secondary antibody for 1h (Cy3 or Alexa-488 anti-mouse or anti-rabbit IgG (1:200)) followed by DAPI for 10 min. The slices were then washed with PBS and mounted with AQUA-MOUNT (Thermo Scientific). Images were taken using a Olympus vs110-ss virtual slide scanner microscope. For the immunohistochemistry on cells, BECs were rinsed in HBSS and fixed in 4% PFA for 15 min at room temperature. The immunostaining was performed using to the same protocol as the brain slices. The cells were mounted and visualized using a Nikon D-Eclipse C1si Spectral Laser confocal microscope. The list of primary antibodies is available in Extended data Table 3.

#### PS2APP Control EC and PS2APP CD2AP EC-KO mice

Animals were deeply anesthetized with isoflurane (0.5 ml/25 g body weight) and transcardially perfused with 20 ml ice-cold PBS. One brain hemisphere was fixed in 4% PFA for 48-72 hr at 4°C with agitation and then transferred to PBS for histopathological analyses. The other hemisphere was sub-dissected into cortical and hippocampal tissues that were frozen and stored at -80°C for biochemical assays. Immersion-fixed hemi-brains were cryoprotected, embedded up to 40 per block in a solid matrix, and coronally sectioned at 35 μm (MultiBrain processing by NeuroScience Associates) as previously described (Kallop et al., 2014; Wang et al., 2011). Sheets of sections were stored in cryoprotectant (30% glycerol, 30% ethylene glycol in PBS) at -20°C until use. Immunohistochemical (IHC) stains for Aβ40 were performed at NeuroScience Associates as described previously (Wang et al., 2011). Primary antibody incubation was followed by three 10 min washes in PBST, followed by incubation with a secondary antibody (donkey anti-goat IgG-Alexa555; 1:500; Thermo Fisher Scientific) for 2 h at room temperature. Tissue sheets were then incubated with Hoechst (1:2500) in PBST (0.1%) for 15 min at room temperature, washed three times (10 min) in PBST (0.1%) and finally three quick washes in PBS. Sheets were mounted onto slides with 0.1% gelatin in PBS and allowed to dry and adhere to the slide at room temperature. Slides were coverslipped with added ProLong Diamond Antifade Mountant (Thermo Fisher Scientific).

All images, segmentation overlays, and data were reviewed by a pathologist. Image acquisition of immunfluorescent slides was performed at 200X magnification using the Nanozoomer S60 or XR (Hamamatsu) digital whole-slide scanner. Ideal exposure for each channel was determined based on samples with the brightest intensity and set for the whole set of slides to run as a batch.

### Cell culture and mouse tissue Western blotting

Cells were lysed directly on plate in RIPA Buffer (50 mM Tris-Cl ph 8.0, 150 mM NaCl, 1% Triton X-100, 0.5% sodium deoxycholate, 0.1% SDS, 1 mM EDTA, 0.2 mM PMSF, cOmplete™ tablet (Roche)) using a cell scrapper. Mouse brain samples were homogenized using a Dounce tissue grinder with ice-cold RIPA buffer. For microvessels extraction: After ice-cold PBS perfusion, and once meninges and white matter were removed, Control EC and CD2AP EC-KO murine brain samples were chopped and frozen in 0.5 mL of MIB containing 0.32 M sucrose and protease and phophatase inhibitors (Bimake) until processed for microvessel enrichment, which was performed as described above for human samples. For both, cells and mouse brain samples, homogenates were sonicated and centrifuged for 15 min at 13,000 rpm. Protein concentration was determined by DC assay (BioRad Laboratories) according to the manufacturer instructions. Proteins were separated using SDS-PAGE and transferred to PVDF membranes (Millipore) for Western blot analysis. Membranes were blocked using 5% skim-milk powder and incubated overnight in primary antibodies. Secondary anti-HRP conjugate antibodies were used and immunoreactivity was visualized using Western LightingTM (PerkinElmer). Immunoreactivity was developed on film or using a ChemiDoc™ imaging system (Biorad), band intensity was quantified using Image Lab and normalized to actin as a loading control. The list of primary antibodies is available in Extended data Table 3.

### A**β** peptide measurements

Frozen hippocampal tissues were homogenized in 10 volumes of TBS (50 mM Tris pH 7.5, 150 mM NaCl) including complete EDTA-free protease inhibitor mixture (Roche Diagnostics) with aprotinin (20 μg/ml) and leupeptin (10 μg/ml) in a QIAGEN TissueLyser II (3 min at 30 Hz). Samples were then centrifuged (10,000 g, 20 min, 4°C; supernatants were collected and stored at -80°C until analyzed. The pellet was homogenized in 10 volumes of 5 M guanidine HCl using the TissueLyser II and then placed on a rotisserie for 3 hr at room temperature. Samples were diluted 1:10 in a casein buffer (0.25% casein/5 mM EDTA, pH 8.0, in PBS), including aprotinin (20 μg/ml) and leupeptin (10 μg/ml), vortexed and centrifuged (20,000 g, 20 min, 4°C). Aβ40 and Aβ42 concentrations in mouse hippocampal samples were measured using ELISA; rabbit polyclonal antibody specific for the C terminus of Aβ40 or Aβ42 (Millipore) was coated onto plates, and biotinylated monoclonal anti Aβ 1-16 (Covance, clone 6E10) was used for detection.

### Immunoprecipitations (IP)

#### From cell lysates

Proteins were extracted from cells as described in the Western blotting section. 1 mg of protein in 500 μl of homogenization buffer (wash buffer with different concentration of ionic and non-ionic detergent, see Figure 3 for detergent content) was added to 10 μl of packed beads: Protein G Sepharose 4 Fast Flow (GE Healthcare Life Sciences) for mouse antibodies or Protein A Sepharose 4 Fast Flow (GE Healthcare Life Sciences) for rabbit antibodies. Sample were inverted at 4°C for 1 hour as a preclear step. At the same time 1 μg of antibody in 500 μl of IP wash buffer (20 mM Tris-Cl ph 7.5, 150 mM NaCl, 0.1% Triton X-100, 0.1% NP40, 0.5 mM EDTA, 0.2 mM PMSF, cOmplete™ tablet (Roche)) was added to 10 μl of packed beads and inverted at 4°C for 1 hour to bind the antibodies to the beads. The tubes were centrifuged to pellet the beads and the antibody solutions were aspirated and replaced with precleared cell lysate and inverted at 4°C for 2 hours to immunoprecipitate the protein of interest. Beads were rinsed 3 times with IP wash buffer and 40 μl of 2x Laemmli Buffer was added to packed beads and boiled for 5 min. at 95°C pelleted and supernatant used for Western blot.

#### From mouse brain homogenates

All steps are identical to the cell lysate IP except that whole mouse brain is homogenized in a Dounce tissue grinder on ice. The homogenate was fractionated into two fractions. The first fraction was generated by homogenizing in brain fraction 1 (BF1) buffer (150 mM NaCl, 1% NP-40, 50 mM Tris-Cl (pH 8.0), cOmplete™ tablet (Roche)) and centrifuged for 15 min. at 12,000g. The supernatant was designated as BF1 and the pellet was resuspended in BF2 buffer (150 mM NaCl, 1% NP-40, 0.5% DOC, 0.1% SDS, 50 mM Tris-Cl (pH 8.0), cOmplete™ tablet (Roche)) and centrifuged at 10,000g and the supernatant designated as BF2.

### Statistical Analysis

Data are presented as mean ± SEM. A ROUT test was used to identify outliers. Details for the statistical analysis for each experiment are described in the figure legends. For all data, when two groups were compared, a Student t-test (paired or unpaired when appropriate) was performed. A repeated measure ANOVA, a two-way ANOVA followed by a Sidak multiple comparisons or a one-way ANOVA with Tukey’s post hoc analysis was utilized when comparing multiple groups. When the variance was not equal, a Welch’s correction was applied. For the cell experiments, protein levels after the CD2AP knockdown were compared to a theoretical mean of 100 using a one-sample t-test. To verify the association between the two parameters, a linear regression was used to generate a correlation coefficient. Values of P<0.05 were accepted as statistically significant. All statistical analyses were performed with Prism 7 (GraphPad software) and SPSS (IBM) softwares.

## Acknowledgments

This work was supported by Canadian Institutes of Health and Research (CIHR, F.C. and M.D.N, grant no: 10021444), a postdoctoral scholarship from CIHR and Alberta Innovates Health solution (M.V.), The Religious Orders Study is supported by the National institute on Aging (P30AG10161, R01AG15819, D.A.B.). Dr Amir Sanati Nezhad for the hCMEC/D3 cell line, Dr Frank Visser for the preparation of the viral vectors, Dr Vincent Emond for carefully proof-reading the manuscript and Dr Justin Chun for his expert opinion on CD2AP EC-KO mice kidney function.

## Author contributions

M.V. designed the study, performed and analyzed the cell culture and awake two photon experiments, analyzed the human data, wrote the manuscript and build the figures, A.I. trained M.V., helped with the experimental design and data analysis for the awake two-photon experiments. B.K. execution of experiments, data analysis and interpretation, revised the manuscript. C.G. set up the protocol for the culture of the primary mouse BECs and helped with the cell culture experiments. S. L. did all the behavioral experiments. J.L. execution of experiments, data analysis. P.B. extracted the protein from the isolated brain vessels, and performed the western blot for the human samples. R.C.M. designed, performed and analyzed the pressure myography experiments. G.P. assisted M.V at the beginning of the two-photon experiments and analyzed vessels spontaneous activity (vasomotion). Y.J. prepared the Reelin solution for the cell culture experiments and did the immunofluorescence on the mouse tissue. S. H. prepared Reelin solution for the awake two-photon imaging. C.B. tested the siRNA for the cell experiment and designed the protocol for the primary BECs cultures. L.R. performed the immunohistochemistry with the human tissue and performed WB on the brain vessels extract of the CD2AP EC-KO mouse line. C.T. extracted proteins from the human samples and performed the ELISA and western blot to characterize the human cortex samples. M. H. helped with the analysis of the MRI images. E.E. Tie2-CD2AP mice kidney analysis. B. M. execution of Tie2-CD2AP-APP mouse-related experiments and data analysis. O. F. pathology work of Tie2-CD2AP-APP mice. M. R-G. generated the Tie2-CD2AP-APP mice. D.M. Supervised E.E. and examined the kidney histology W.N. wrote the script for the analysis of the awake two-photon experiments. J. K. provided the AAVBR1 virus. J. D. Supervised M.H. and reviewed the manuscript. A.P.B. supervised R.C.M. for the pressure myography experiments. D.A.B., provided the human samples and clinical data from the Religious Orders Study participants, and critically reviewed the manuscript. G.R.G. supervised G.P. and A.I. for the awake two-photon experiments. A.S. provided the Tie2-CD2AP mouse lines and critically revised the manuscript. F.C. provided the human data and supervised P.B., L.R. and C.T. for the human experiments. M.D.N. supervised M.V. C.G., Y.J. S.H. S.L. and C.B., designed the experiments and wrote the manuscript.

