## Supplemental file for "The Alzheimer risk factor CD2AP causes dysfunction of the brain vascular network"

**Table S1: Characteristics of participants from the Religious Order Study**

| Characteristics | NCI | MCI | AD | Statistical Analysis |
|---|---|---|---|---|
| *N* | 20 | 20 | 20 | ___ |
| Men, % | 20 | 45 | 35 | C; Pearson test, $\chi 2 = 2.85$; $p = 0.24$ |
| Mean age at death | 87.1 (5.8) | 87.1 (5.2) | 87.3 (4.9) | A; $F(groups)_{2,57} = 0.01$; $p = 0.99$ |
| Mean education, years | 18.5 (3.6) | 18.6 (3.0) | 17.5 (3.0) | A; $F(groups)_{2,57} = 0.69$; $p = 0.51$ |
| Mean MMSE | 27.2 (1.8) | 25.5 (3.1) | 15.8 (7.9)¶ | A; $F(groups)_{2,57} = 30.17$; $p < 0.0001$ |
| Global cognition score | -0.03 (0.38) | -0.43 (0.45) | -1.66 (0.89)¶ | A; $F(groups)_{2,56} = 38.03$; $p < 0.0001$ |
| *APOE4* allele carriage (%) | 30 | 30 | 35 | C; Pearson test, $\chi 2 = 0.15$; $p = 0.93$ |
| Reagan score 3/2/1 (*n*) | 12/8/0 | 10/9/1 | 4/7/9 | ___ |
| CERAD score 4/3/2/1 (*n*) | 7/3/8/2 | 7/3/6/4 | 1/1/6/12 | ___ |
| Cerebellar pH | 6.35 (0.34) | 6.32 (0.28) | 6.31 (0.46) | A; $F(groups)_{2,57} = 0.07$; $p = 0.93$ |
| Postmortem delay (hours) | 7.1 (5.6) | 7.8 (5.2) | 7.8 (4.8) | A; $F(groups)_{2,57} = 0.13$; $p = 0.88$ |
| Neuron Diffuse Plaque Counts | 8.7 (15.9) | 14.5 (13.2) | 19.6 (18.2) | A; $F(groups)_{2,57} = 2.36$; $p = 0.10$ |
| Neuron Plaque Counts | 6.3 (8.9) | 7.0 (7.7) | 18.0 (15.5)* | A; $F(groups)_{2,57} = 6.81$; $p = 0.0022$ |
| Neurofibrillary Tangle Counts | 0.15 (0.49) | 0.25 (0.72) | 5.25 (11.10)& | A; $F(groups)_{2,57} = 4.12$; $p = 0.02$ |
| Soluble Aβ40 concentration (fg/µg protein) | 428.9 (725.7) | 322.8 (649.3) | 340.1 (527.6) | A; $F(groups)_{2,57} = 0.16$; $p = 0.86$ |
| Soluble Aβ42 concentration (fg/µg protein) | 1232 (1220) | 1240 (1358) | 1872 (1450) | A; $F(groups)_{2,57} = 1.49$; $p = 0.23$ |
| Insoluble Aβ40 concentration (pg/mg tissue) | 2749 (3697) | 1094 (1931) | 2224 (3542) | A; $F(groups)_{2,38} = 0.81$; $p = 0.45$ |
| Insoluble Aβ42 concentration (pg/mg tissue) | 622.9 (646.7) | 678.6 (638.6) | 1204 (788.0)& | A; $F(groups)_{2,50} = 3.74$; $p = 0.03$ |

Soluble and insoluble Aβ levels were quantified by ELISA in parietal cortex samples and are presented as a log. ¶ $p < 0.0001$ vs NCI et vs MCI. * $p < 0.001$ vs NCI et vs MCI. & $p < 0.05$ vs NCI et vs MCI. A = ANOVA. C = contingency. NCI: no cognitive impairment. MCI: mild cognitive impairment. MMSE: mini mental state examination. APOE4: apolipoprotein E epsilon 4. CERAD: consortium for Consortium to Establish a Registry for Alzheimer's Disease.

**Table S2: Correlations between CD2AP levels in the parietal cortex and isolated cerebrovascular extracts and AD hallmarks**

| Parameter | CD2AP in whole homogenates of parietal cortex samples | | CD2AP in isolated brain vessels of parietal cortex samples | |
|---|---|---|---|---|
| **Amyloid pathology in whole homogenates of parietal cortex** | *r* | *p* | *r* | *p* |
| ***Neuritic plaque counts*** | -0.021 | 0.873 | -0.123 | 0.368 |
| ***Diffuse plaque counts*** | -0.001 | 0.994 | 0.1 | 0.463 |
| ***Soluble Aβ42 (log)*** | -0.307 | 0.017* | -0.126 | 0.354 |
| ***Soluble Aβ40 (log)*** | -0.261 | 0.046* | -0.165 | 0.229 |
| ***Insoluble Aβ42 (log)*** | -0.076 | 0.582* | -0.097 | 0.497 |
| ***Insoluble Aβ40 (log)*** | -0.272 | 0.081 | -0.176 | 0.283 |
| **Tau pathology in whole homogenates of parietal cortex** | | | | |
| ***Neurofibrillary tangle counts*** | 0.018 | 0.893 | -0.083 | 0.542 |
| ***Total insoluble tau (Tau13)*** | -0.504 | 0.00004* | -0.084 | 0.537 |
| ***Paired helical filament tau (AD2)*** | -0.13 | 0.323 | -0.14 | 0.305 |
| ***Paired helical filament tau (PHF1)*** | -0.166 | 0.205 | -0.145 | 0.285 |
| **Cognition** | | | | |
| ***Last MMSE*** | -0.01 | 0.939 | 0.296 | 0.027* |
| **Global cognitive score** | 0.028 | 0.832 | 0.283 | 0.036* |
| ***Episodic memory*** | 0.036 | 0.789 | 0.269 | 0.047* |
| ***Semantic memory*** | -0.011 | 0.934 | 0.25 | 0.066 |
| ***Working memory*** | 0.06 | 0.65 | 0.135 | 0.327 |
| ***Perceptual speed*** | 0.007 | 0.956 | 0.326 | 0.015* |
| ***Visuospatial memory*** | -0.192 | 0.145 | 0.046 | 0.738 |

Soluble and insoluble levels of Aβ were quantified by ELISA in samples from human parietal cortex. The correlations were evaluated using a linear regression analysis. Significant correlations (P<0.05) are highlighted in blue. *Correlation still significant after adjustment for age of death and education. MMSE: mini mental state examination.

### Table S3: List of primary antibodies

| Antibody | Clone | Specificity | Host | Source |
|---|---|---|---|---|
| Actin | C4 | a.a. 50-70 | Mouse | Millipore |
| pAkt Ser473 | Polyclonal | Endogenous levels of Akt1 when phosphorylated at Ser473 | Rabbit | Cell signaling |
| Akt 1-2-3 | Polyclonal | Endogenous levels of total Akt1, Akt2 and Akt3 proteins | Rabbit | Cell signaling |
| ApoER2 Full length | EPR3326 | Human ApoER2 aa 950 to the C-terminus (C terminal) | Rabbit | Abcam |
| ApoER2 45KD | polyclonal | Human ApoER2 internal sequence amino acids 396-445 | Rabbit | Abcam |
| CD2AP | Polyclonal | n/a | Rabbit | Protein Tech |
| CD2AP | Polyclonal | n/a | Rabbit | Sigma-Aldrich |
| Claudin 5 | clone 4C3C2 | Synthetic peptide derived from the mouse Claudin-5 protein | Mouse | ThermoFisher Scientific |
| Dynamin II | polyclonal | Human Dynamin 2 a.a. 760-779. | Rabbit | BD Biosciences |
| GFP | B-2 | aa 1-238 representing full length GFP (green fluorescent protein) of Aequorea victoria origin | Mouse | Santa Cruz |
| Laminin | Polyclonal | Pan-specific | Rabbit | Novus Biological |
| NeuN | clone EPR12763 | Human NeuN aa 1-100 (Cysteine residue) | Rabbit | Abcam |
| PICALM | Polyclonal | C-terminus of CALM of human origin | Goat | Santa Cruz |
| Rabaptin 5 | Polyclonal | Human Rabaptin-5 a.a. 247-417. | Mouse | BD Biosciences |
| Reelin | G10 | Recombinant fusion protein, corresponding to amino acids 164-496 of mouse Reelin | Mouse | Abcam |
| Reelin | RE-3B9(R3B9) | Reln CR50 | Mouse | MBL |
| Type IV Collagen | Polyclonal | Human and bovine placental collagen type IV | Goat | Millipore |
| VE-Cadherin | Polyclonal | C-terminus of VE-cadherin of human origin | Goat | Santa Cruz |

**Table S4: DNA sequences of the shRNAs targeting mouse CD2ap (accession NM_009847) in each AAV construct**

| Construct | 97-mer shRNA sequence |
|---|---|
| pAAV-U6-Dh09shRNA-CMV-GFP-WPRE-hgHpA or pAAV-U6-Dh09shRNA-tdTomato-GFP-WPRE-hgHpA | TGCTGTTGACAGTGAGCGACCATCAAACGGGAAAGGCAAGTAGTGAAGCCACAGATGTACTTGCCTTTCCCGTTTGATGGGTGCCTACTGCCTCGGA |
| pAAV-U6-Dh10shRNA-CMV-GFP-WPRE-hgHpA or pAAV-U6-Dh10shRNA-tdTomato-GFP-WPRE-hgHpA | TGCTGTTGACAGTGAGCGCAGGGTGAACTGAACGGTAAAGTAGTGAAGCCACAGATGTACTTTACCGTTCAGTTCACCCTTTGCCTACTGCCTCGGA |
| pAAV-U6-Dh11shRNA-CMV-GFP-WPRE-hgHpA or pAAV-U6-Dh11shRNA-CMV-tdTomato-WPRE-hgHpA | TGCTGTTGACAGTGAGCGAGCGGTATGGCTACAGACCAAGTAGTGAAGCCACAGATGTACTTGGTCTGTAGCCATACCGCCTGCCTACTGCCTCGGA |
| pAAV-U6-Dh12shRNA-CMV-GFP-WPRE-hgHpA or pAAV-U6-Dh12shRNA-CMV-tdTomato-WPRE-hgHpA | TGCTGTTGACAGTGAGCGCAAGTCAGATTTCTTGTGTTATTAGTGAAGCCACAGATGTAATAACACAAGAAATCTGACTTTTGCCTACTGCCTCGGA |
| pAAV-U6-scAshRNA-CMV-GFP-WPRE-hgHpA or pAAV-U6-scAshRNA-tdTomato-GFP-WPRE-hgHpA | TGCTGTTGACAGTGAGCGCGCCAGTATTTGCCCTTGATATTAGTGAAGCCACAGATGTAATATCAAGGGCAAATACTGGCATGCCTACTGCCTCGGA |
| pAAV-U6-scBshRNA-CMV-GFP-WPRE-hgHpA or pAAV-U6-scBshRNA-CMV-tdTomato-WPRE-hgHpA | TGCTGTTGACAGTGAGCGAACCACTAAGCTTGCTTCTTACTAGTGAAGCCACAGATGTAGTAAGAAGCAAGCTTAGTGGTCTGCCTACTGCCTCGGA |
| pAAV-U6-scCshRNA-CMV-GFP-WPRE-hgHpA or pAAV-U6-scCshRNA-CMV-tdTomato-WPRE-hgHpA | TGCTGTTGACAGTGAGCGATGTACAGGACTTCACAGAAAATAGTGAAGCCACAGATGTATTTTCTGTGAAGTCCTGTACAGTGCCTACTGCCTCGGA |
| pAAV-U6-scramb-CMV-GFP-WPRE-hgHpA or pAAV-U6-scramb-CMV-tdTomato-WPRE-hgHpA | TGCTGTTGACAGTGAGCGAAACAGCAAGTAACGCAGACGCTAGTGAAGCCACAGATGTAGCGTCTGCGTTACTTGCTGTTCTGCCTACTGCCTCGGA |

Antisense sequences are underlined

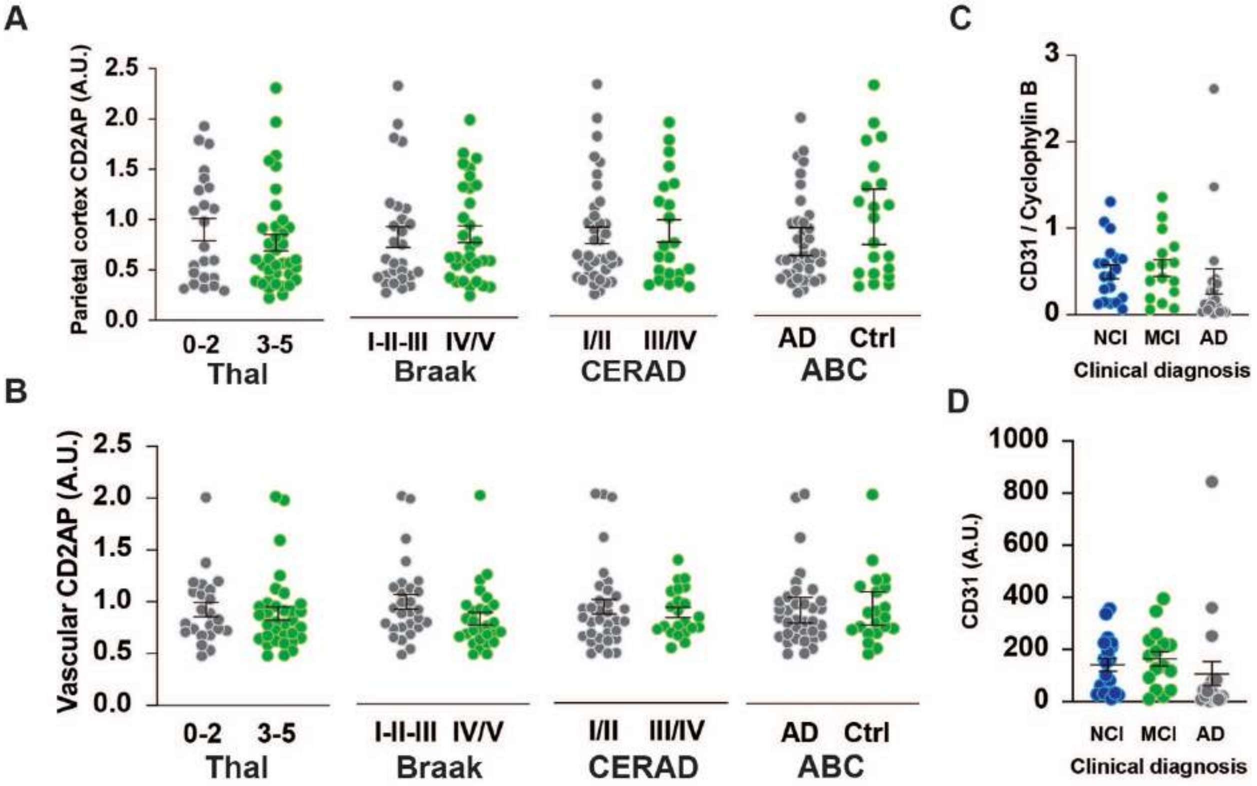


**Figure S1: CD2AP levels in the cortex and isolated brain vessels of AD volunteers.**
A) CD2AP levels in parietal cortex samples from volunteers classified according to Thal (0-2: n = 23, 3-5: n = 3, Unpaired Student t-test), Braak (I-II-III: n = 32, IV/V: n = 28, Unpaired Student t-test), CERAD (I/II: n = 38, III/IV: n = 22, Unpaired Student t-test) or ABC (Control: n = 21, AD: n = 39, Unpaired Student t-test) scores. B) CD2AP levels in isolated brain microvessels of volunteers classified according to Thal (0-2: n = 23, 3-5: n = 33, Unpaired Student t-test), Braak (I-II-III: n = 28, IV/V: n = 28, Unpaired Student t-test), CERAD (I/II: n = 34, III/IV: n = 22, Unpaired Student t-test with Welch's correction) or ABC (Control: n = 21, AD: n = 35, Unpaired Student t-test) scores. Levels of the endothelial cell marker CD31 in the isolated brain vessels from volunteers classified according to the clinical diagnosis (NCI, n=19, MCI, n=16, AD, n=19, One-way ANOVA) C) Normalized on cyclophilin B or D) not normalized.
Brain samples were obtained from participants in the Religious Orders Study, a longitudinal study of aging and dementia with an extensive amount of clinical and neuropathological data [1]. Values for CD31 levels can also be found in [2].
CD2AP: CD2-associated protein.

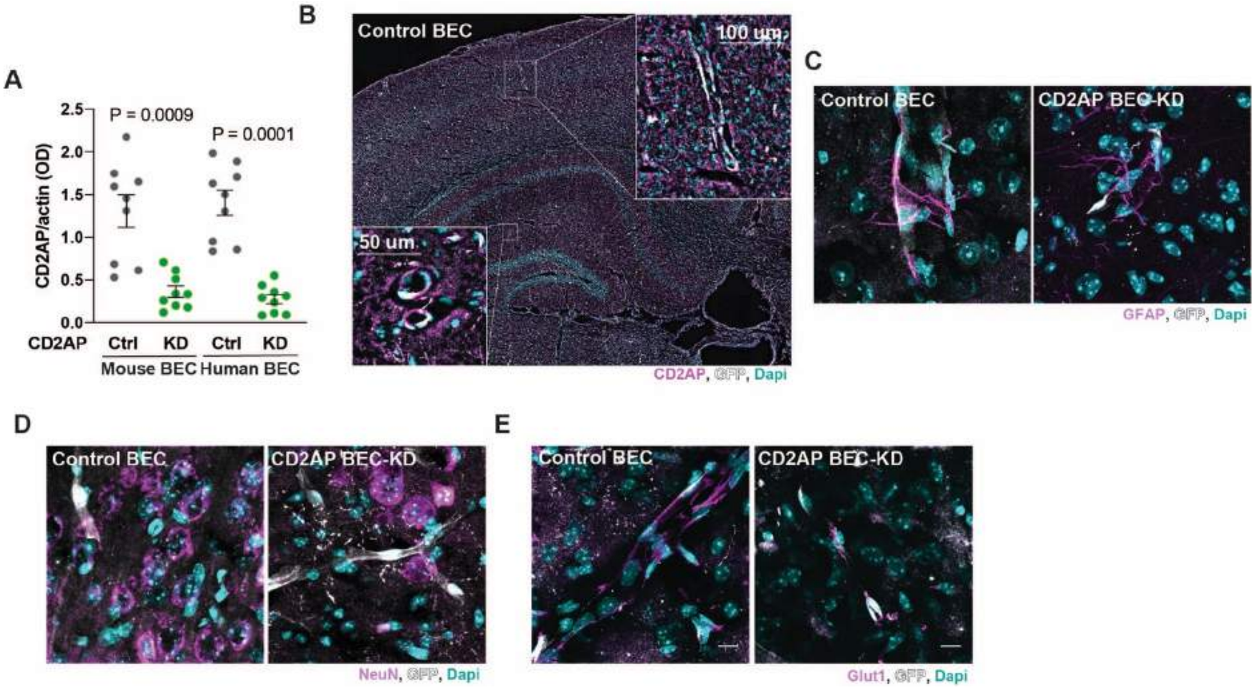


**Figure S2: AAVBR1 targets brain endothelial cells in mice**
A) Level of CD2AP in mouse (m) primary brain endothelial cells and hCMEC/D3 immortalized cell line (h) after transfection with CD2AP siRNA. N = 9 wells per group. mControl and mCD2AP KD and hControl and hCD2AP KD were compared using an Unpaired Student t-test with a Welch's correction. Immunofluorescent image of brain slice from Control BEC and CD2AP BEC-KD mice stained for B) CD2AP (magenta), GFP (white) and Dapi (cyan), C) GFAP (magenta), GFP (white) and Dapi (cyan), D) NeuN (magenta), GFP (white) and Dapi (cyan), E) Glut1 (magenta), GFP (white) and Dapi (cyan).

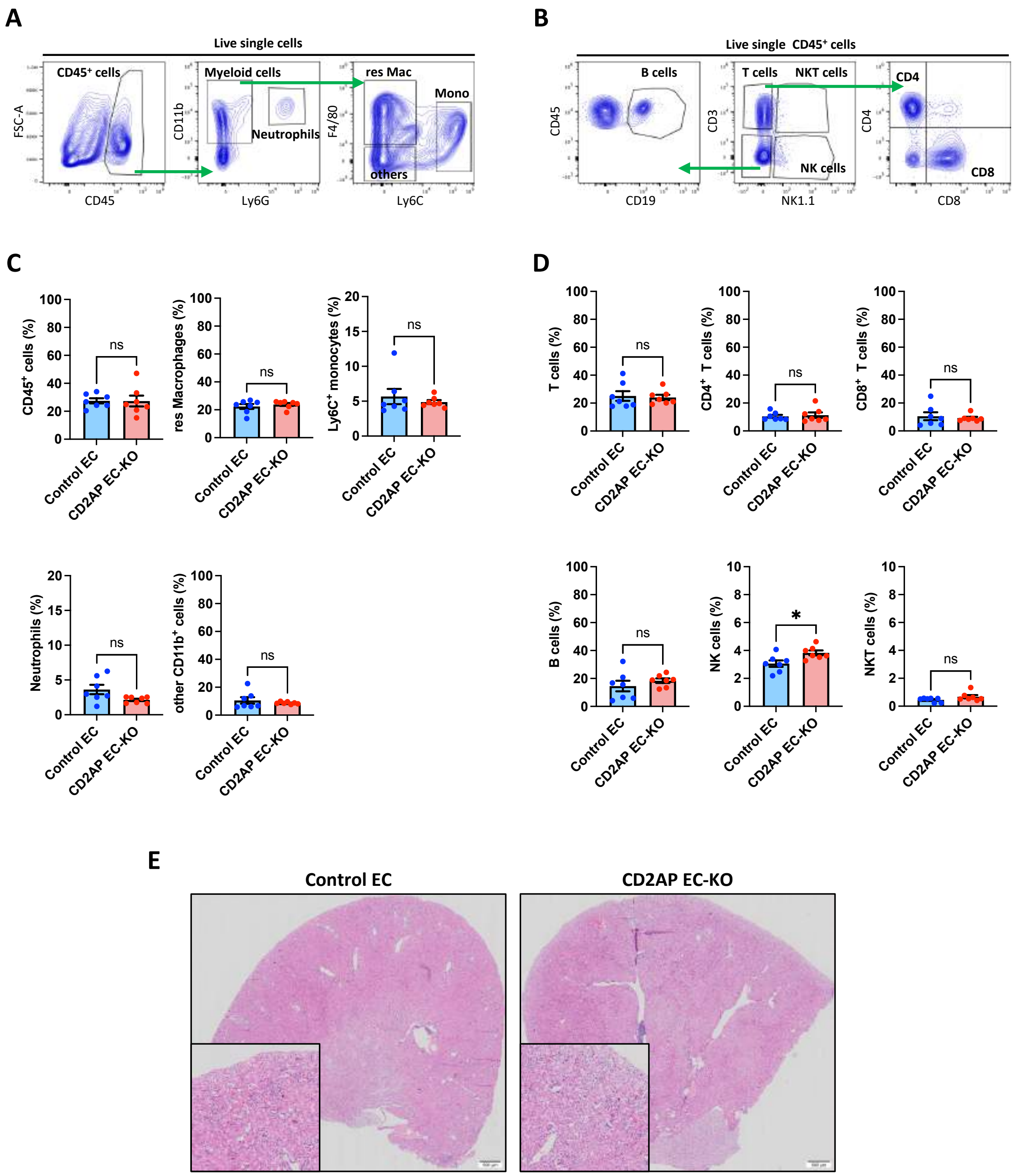


**The absence of CD2AP does not alter the renal immune environment or structure.** Kidneys from control EC and CD2AP EC-KO mice (age: 30-32 week) were harvested and used for multiparameter flow cytometry to study the renal immune composition and for histopathology. **(A)** Gating strategy used to study the myeloid cell subset of immune cells in the kidney. $CD45^+$ cells were selected from live single cells, then neutrophils ($CD45^+$ $CD11b^+$ $Ly6G^+$ cells), resident macrophages (res Mac or $CD45^+CD11b^+F4/80^+Ly6G^-$ cells), $Ly6C^+$ monocytes (Mono or $CD45^+CD11b^+Ly6C^+Ly6G^-$ cells) and other myeloid cells (others or $CD45^+CD11b^+Ly6C^-F4/80^-Ly6G^-$ cells) were selected from $CD45^+$ cells. **(B)** Gating strategy used to study the lymphoid cell subset in the kidney. $CD45^+$ cells were selected from live single cells, then T cells ($CD45^+CD3^+NK1.1^-$ cells), $CD4^+$ T cells ($CD45^+CD3^+CD4^+NK1.1^-$ cells), $CD8^+$ T cells ($CD45^+CD3^+CD8^+NK1.1^-$ cells), B cells ($D45^+CD19^+CD3^-NK1.1^-$ cells), NK cells

(D45$^+$NK1.1$^+$CD3$^-$CD19$^-$ cells) and NKT cells (D45$^+$NK1.1$^+$CD3$^+$CD19$^-$ cells) were selected from CD45$^+$ cells. **(C)** Quantification of total leukocytes (CD45$^+$ cells), resident macrophages, Ly6C$^+$ monocytes, neutrophils, and other myeloid cells. The results are presented as the mean ± SEM. An unpaired t test was performed, where * $p<0.05$, ns= not significant (N=7/condition). **(D)** Quantification of total T cells, CD4$^+$ T cells, CD8$^+$ T cells, B cells, NK cells and NKT cells. The results are presented as the mean ± SEM. An unpaired t test was performed, where ns= not significant (N=7/condition). **(E)** Hematoxylin and Eosin staining of kidneys from control EC and CD2AP EC-KO mice showing no gross alterations in renal morphology (N=3/condition).

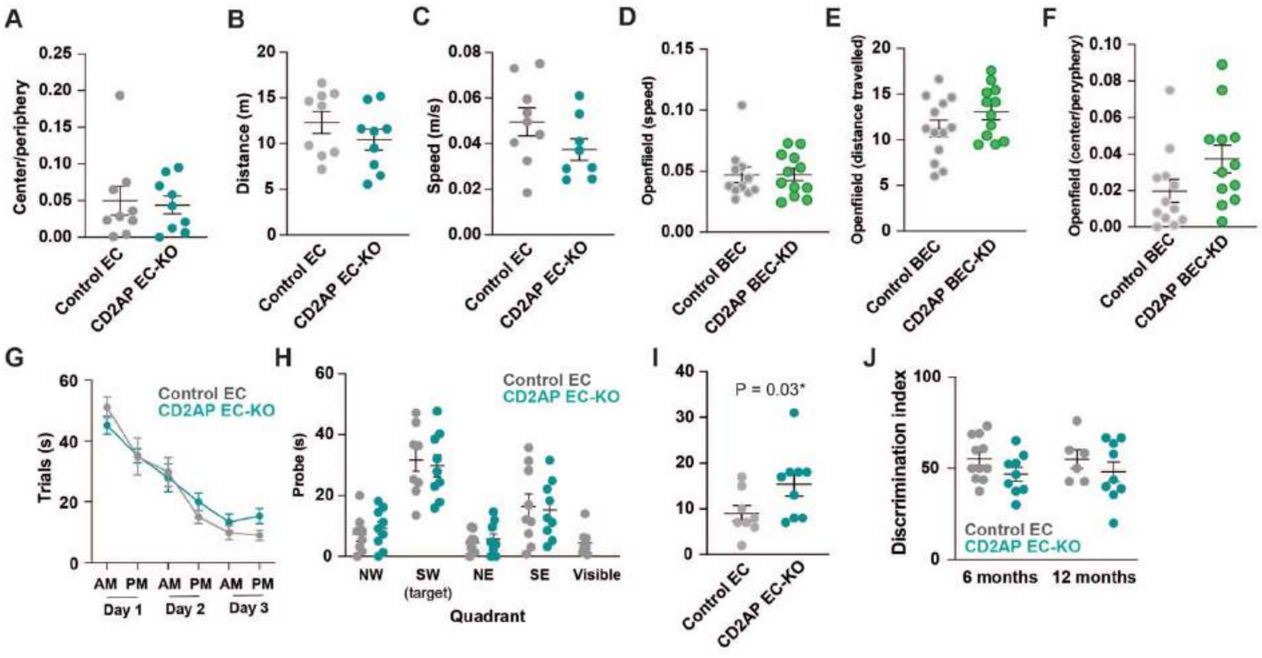


**Figure S4: Behavioral consequences of endothelial-CD2AP downregulation in mice**
Control EC and CD2AP EC-KO A) Center/periphery ratio, B) Distance travelled and C) Speed during the openfield.
Control BEC and CD2AP BEC-KD D) Center/periphery ratio, E) Distance travelled and F) Speed during the openfield.
Time to reach the reach the target during the G) trials and the H) probe test and time to target in at I) Day3 PM during the Morris Water Maze in Control EC and CD2AP EC-KO mice.
J) Discrimination index during the novel object recognition test in Control EC and CD2AP EC-KO mice.

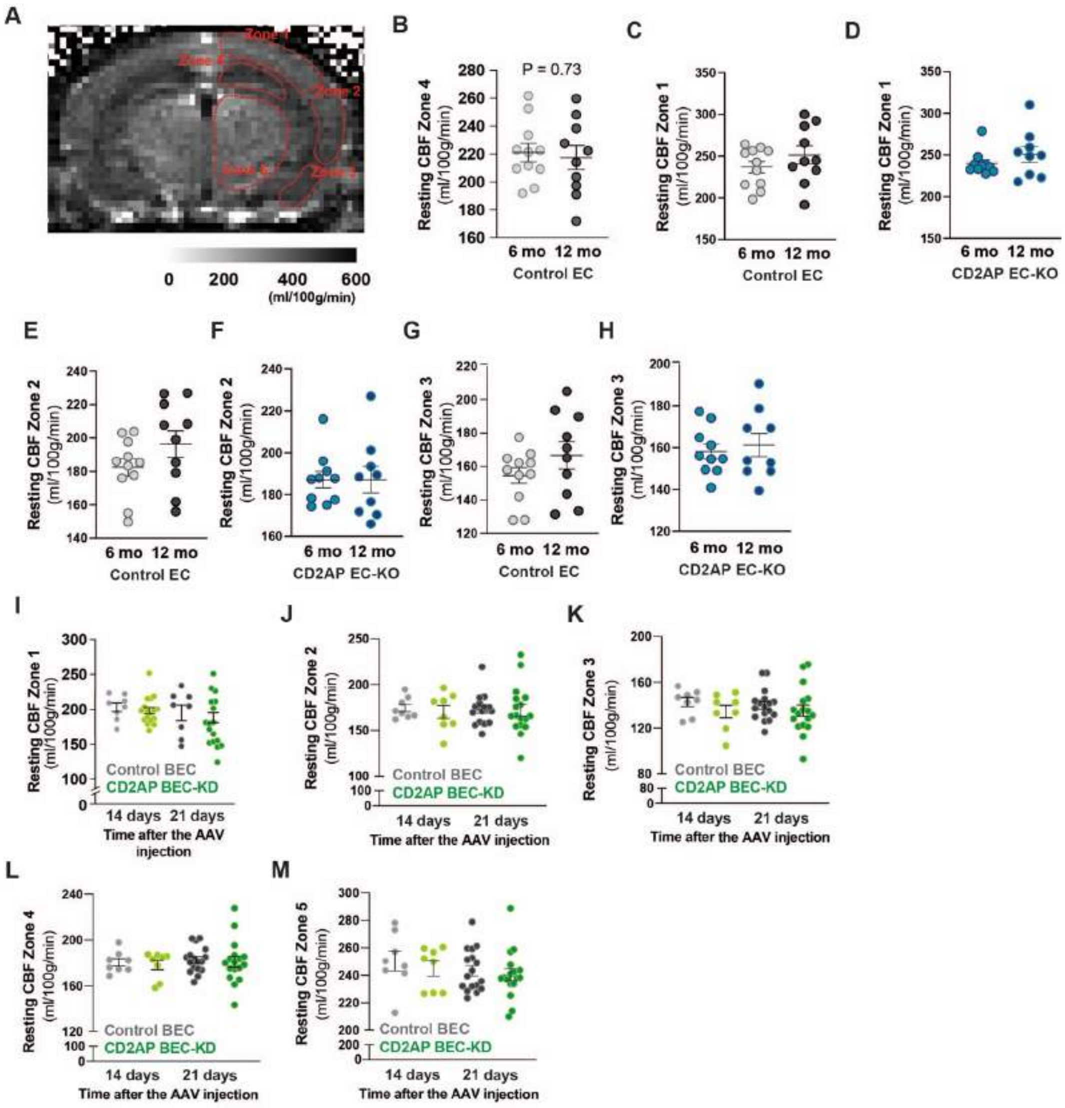


**Figure S5: Resting cerebral blood flow in mice with endothelial CD2AP downregulation.**
A) Identification of the zones where the measurement of cerebral blood flow were done. B to H) Measurements for Control EC and CD2AP EC-KO mice.
I to M) Measurements for Control BEC and CD2AP BEC-KO mice.
Data are presented as mean and SEM.

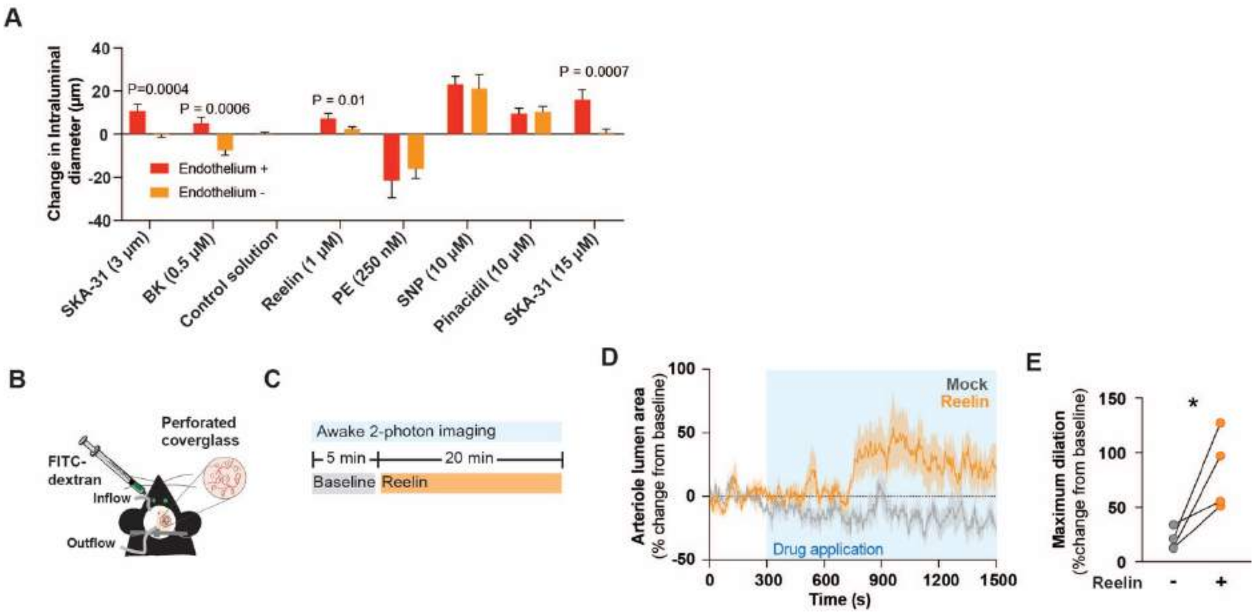


**Figure S6: Reelin increases brain vessels dilation**

A) Intraluminal diameter changes in single cannulated cerebral arteries from 5-6 weeks old mice (n = 4-8) with endothelium and without endothelium following generation of myogenic tone at 70 mmHg, arteries were bath exposed to SKA-31 (3 and 15 μM), bradykinin (BK, 0.5 μM), control solution (volume matched with Reelin), Reelin (1 μM), and the endothelium-independent agent phenylephrine (PE, 250 nM), sodium nitroprusside (SNP, 10 μM) and pinacidil (10 μM) in the presence and absence of endothelium. Maximal passive vessel diameter at the end of the protocol was obtained in the presence of Krebs' buffer containing 2 mM EGTA and zero added $CaCl_2$. Unpaired Student t-test. B) Schematic of the cranial window with the perforated coverslip. C) Timeline of the drug application during the experiment. D) Effect of control solution (Mock) or Reelin (1 μM) on penetrating arterioles transversal luminal area (n = 4 mice). E) Maximum dilation of a penetrating arteriole in response to control solution or Reelin (n = 4 mice). Each dot represents one mouse. *Paired Student t-test.

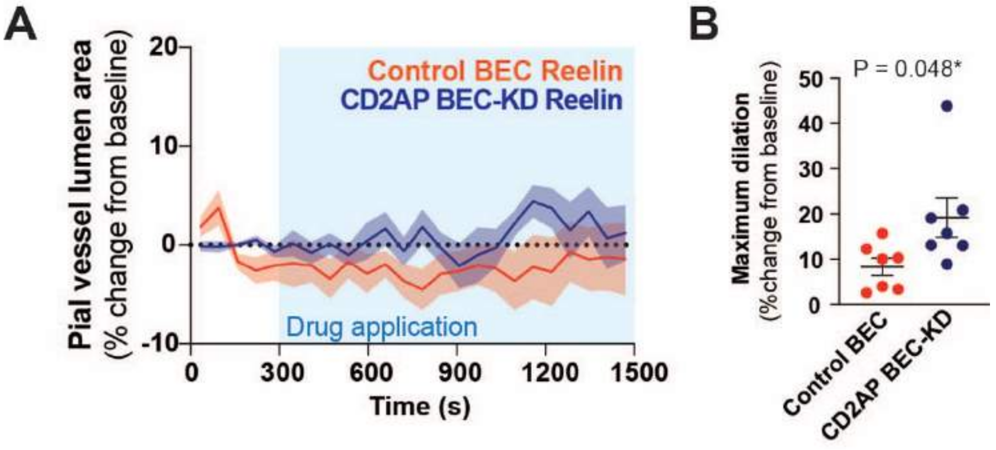


**Figure S7: Control BEC and CD2AP BEC-KD mice pial vessels response to Reelin superfusion.**
A) Effects of Control BEC and CD2AP BEC-KD on pial vessel luminal area in response to Reelin (1 μM) superfusion. B) Maximal dilation of pial vessels in response to Reelin (Unpaired Student t-test, each dot represents one animal).
Data are presented as mean ± SEM.

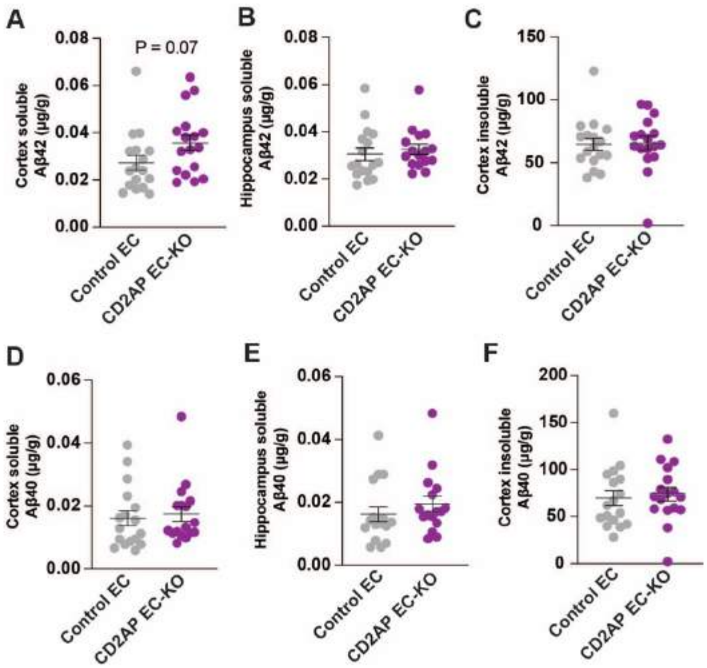


**Figure S8: Endothelial CD2AP loss impact on Aβ accumulation in the PS2APP mouse model of AD.**

Levels of soluble Aβ42 in the A) cortex and B) hippocampus.

C) Levels of insoluble Aβ42 in the hippocampus.

Levels of soluble Aβ40 in the D) cortex and E) hippocampus.

F) Levels of insoluble Aβ40 in the hippocampus.

Data were compared using unpaired Student t-test. No difference was found between groups.

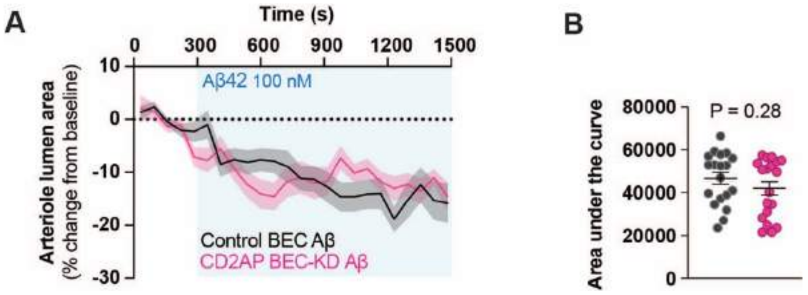


**Figure S9: Aβ application in Control and CD2AP BEC-KD mice**

A and B) Control BEC (black) and CD2AP BEC-KD (pink) arteriole lumen area change with Aβ42 (100 nM) application.

Data are presented as mean and SEM.

Data were compared using an Unpaired Student t-test. No difference was found.

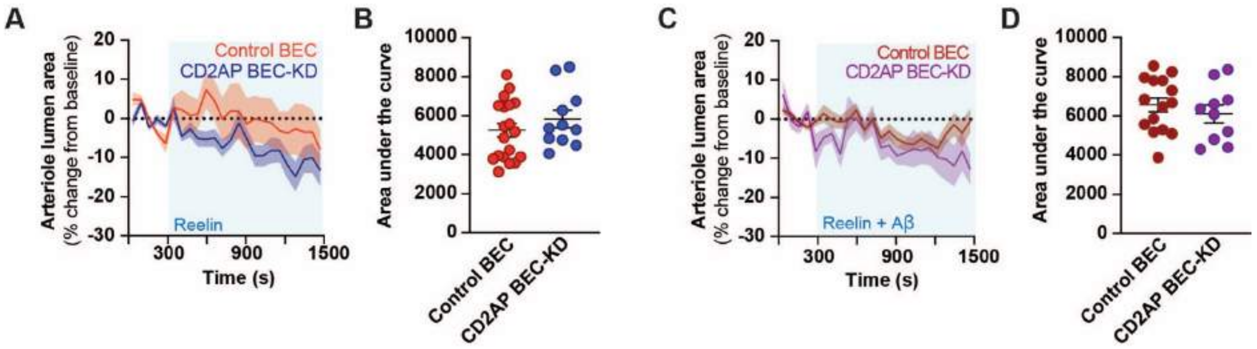


**Figure S10: Drug application in Control and CD2AP BEC-KD mice**

A and B) Control BEC (orange) and CD2AP BEC-KD (blue) arteriole lumen area change with Reelin (1 $\mu$M) application.

C and D) Control BEC (brown) and CD2AP BEC-KD (purple) arteriole lumen area change with Reelin (1 $\mu$M) + A$\beta$42 (100 nM) application.

Data are presented as mean and SEM.

Data were compared using an Unpaired Student t-test. No difference was found.

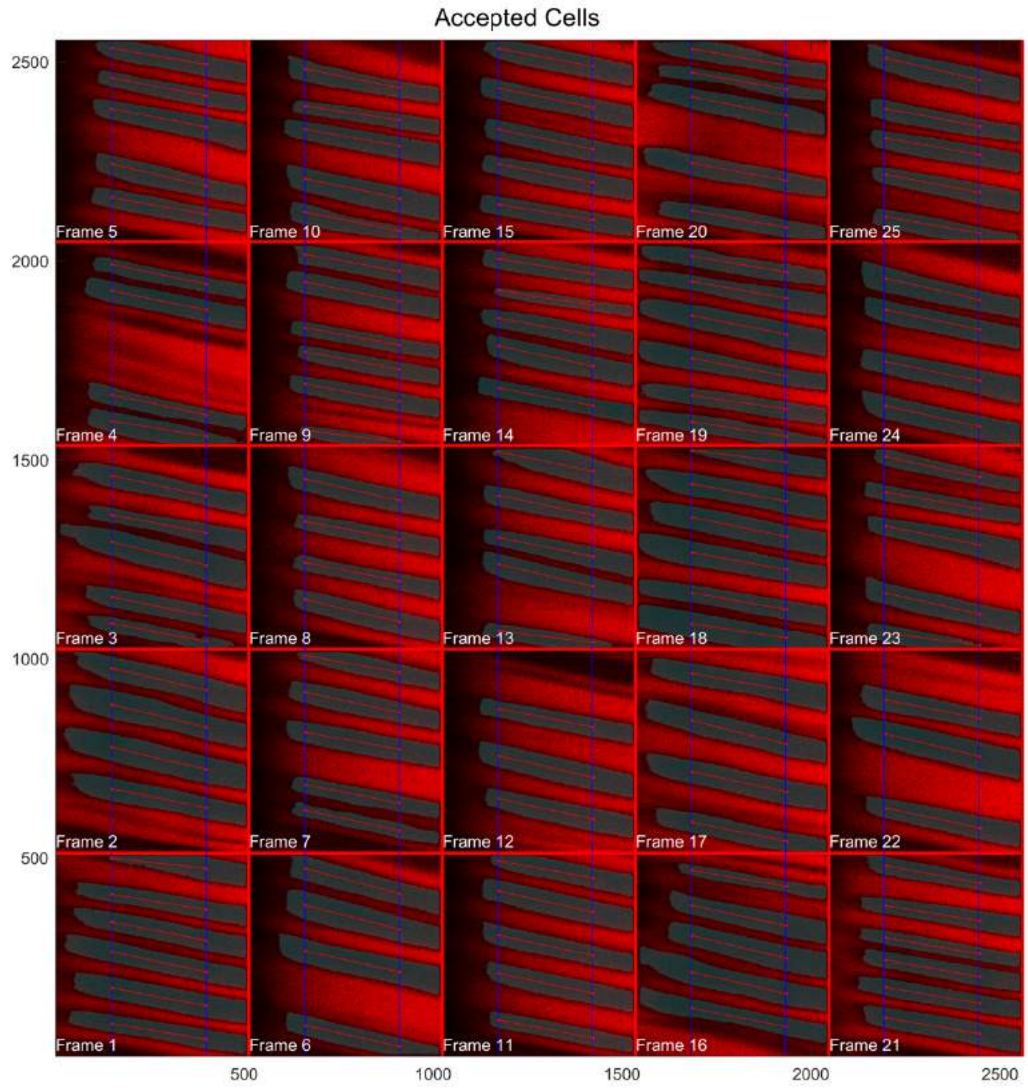


**Figure S11: Ouput example from the capillary red blood cells analysis for U-Net slow. Gray: Red blood cells picked up by U-Net slow.**

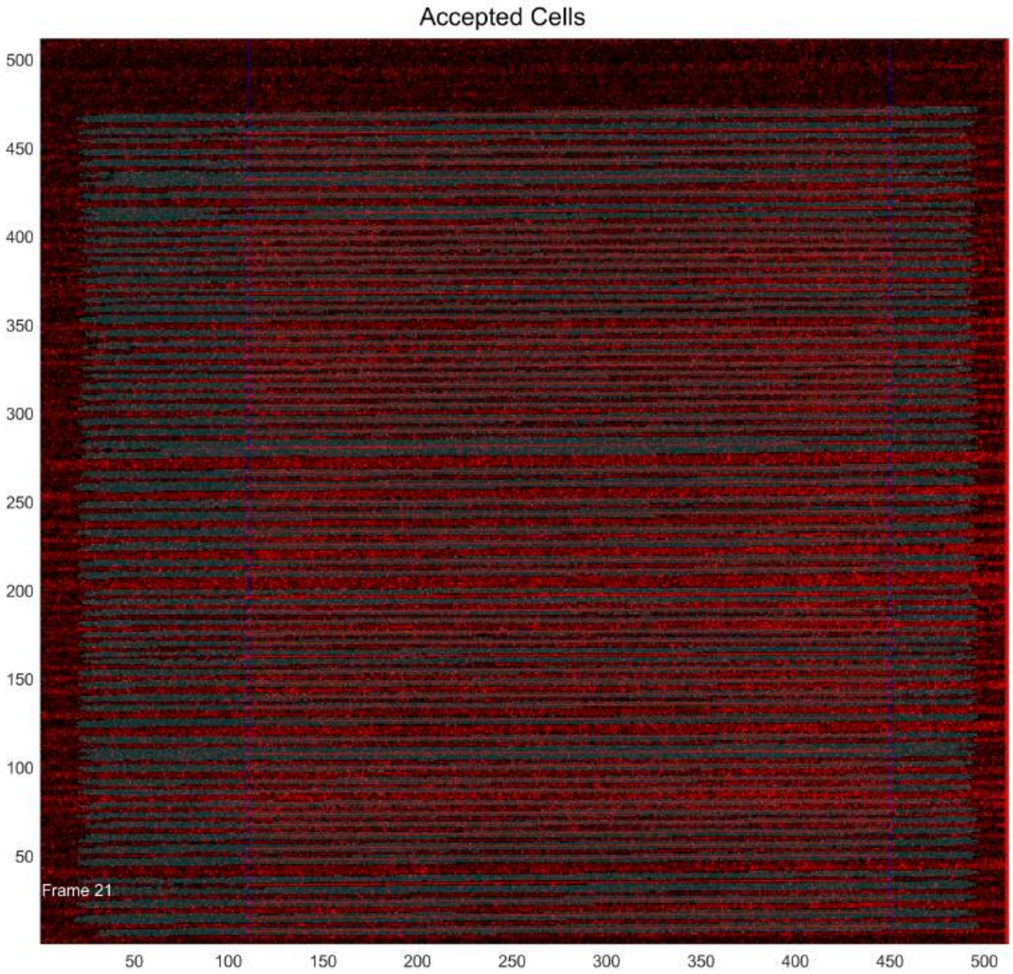


**Figure S12: Ouput example from the capillary red blood cells analysis for U-Net fast. Gray: Red blood cells picked up by U-Net fast.**

### Supplementary materials and methods

#### Reagents

The following fluorophore-conjugated anti-mouse antibodies were purchased from BioLegend (San Diego, CA, USA): BV605-CD11b (clone M1/70), FITC-CD45 (clone 30-F11), PerCP-Ly6G (clone 1A8), PE-F4/80 (clone BM8), PE/Cy7-Ly6C (clone HK1.4), PerCP/Cy5.5-NK1.1 (clone PK136), BV421-CD19 (clone 6D5), PE-CD3e (clone KT3.1.1), PE/Cy7-CD4 (clone GK1.5). The fluorophore-conjugated anti-mouse APC/Cy7-CD8 (clone 53-6.7) antibody was purchased from BD Bioscience (Franklin Lakes, New Jersey, USA). Multi Tissue Dissociation Kit 1 was purchased from Miltenyi Biotec.

#### Method

#### Flow Cytometry.

Murine kidney tissue was isolated from mice after perfusion via saline injection into the heart. Kidney capsule and fat were removed before incubation of whole tissue in Multi Tissue Dissociation Kit 1 following the manufacturer's instructions. Tissue homogenate was then passed through a 40- μM nylon cell strainer and washed with cold PBS (phosphate buffered saline). Cell pellet was incubated in ACK lysis buffer (Gibco) for 3 minutes to remove red blood cells and washed in PBS. Viable cells were isolated using a density gradient by mixing cells with Percoll (GE Healthcare) and centrifuging at 800 g for 25 minutes. Cells were washed in staining buffer (PBS, EDTA 2 mM, BSA 1%), and labeled with the following anti-mouse antibodies: FITC-CD45 (clone 30-F11), BV605-CD11b (clone M1/70), PerCP-Ly6G (clone 1A8), PE-F4/80 (clone BM8), PE/Cy7-Ly6C (clone HK1.4), PerCP/Cy5.5-NK1.1 (clone PK136), BV421-CD19 (clone 6D5), PE-CD3e (clone KT3.1.1), PE/Cy7-CD4 (clone GK1.5) and/or APC/Cy7-CD8 (clone 53-6.7). Cells were resuspended in staining buffer before analysis by a Attune NxT (Life Technologies) and FlowJo software (BD Biosciences).

#### Hematoxylin & Eosin staining.

Murine kidney was fixed in 10% neutral buffered formalin, followed by dehydrating in gradual ethanol solution and embedding with paraffin. Subsequently, the embedded tissues were cut into 5 µm serial sections, which were following dewaxed in xylene and rehydrated in ethanol gradients. Ultimately, tissues were stained with hematoxylin-eosin (H&E) for histological evaluation, and sections were captured by using a light microscope (Olympus) with a 20x magnification.
